# Extracellular Stimulation and Ephaptic Coupling of Neurons in a Fully Coupled Finite Element-Based Extracellular–Membrane–Intracellular (EMI) Model

**DOI:** 10.1101/2025.11.27.690917

**Authors:** Karoline Horgmo Jæger, Aslak Tveito

**Affiliations:** Simula Research Laboratory, Norway

## Abstract

The extracellular potential surrounding neurons is of great importance: it is measured to interpret neural activity, it underpins ephaptic coupling between neighboring cells, and it forms the basis for external stimulation of neural tissue. These phenomena have been studied for decades, both experimentally and computationally. In computational models, variants of the classical cable equation for membrane dynamics and an electrostatic equation for the extracellular field are the most common approaches. Such formulations, however, typically decouple the governing equations and therefore neglect the biophysical coupling between the extracellular (E) space, the cell membrane (M), and the intracellular (I) space. Here, we use a finite element–based Extracellular–Membrane–Intracellular (EMI) approach that solves a fully coupled system of equations to study extracellular stimulation and ephaptic coupling of cerebellar Purkinje neurons and neocortical layer 5 pyramidal neurons. These two archetypes differ substantially in morphology, ion-channel distribution, and firing behavior, and together span a range of neuronal properties. Specifically, we assess responses to extracellular stimulation while varying the distance to the stimulation source, the amplitude, and the frequency of the external current. We also investigate ephaptic interactions between neurons, and examine how the firing pattern of one neuron can be affected by the firing pattern of a neighboring neuron, how the rate of synchronization between neighboring neurons depends on cell distance and extracellular conductivity, and finally whether a neuron can directly trigger excitation in a neighboring neuron through ephaptic coupling. The results provide quantitative insight into extracellular field–mediated neural coupling and how externally applied fields, such as those used in deep brain stimulation, interact with single-neuron biophysics.

## 1 Introduction

The extracellular potential (EP) is of fundamental importance for investigating the properties and dynamics of neural tissue [1, 2]. It is also essential for understanding the biophysical mechanisms underlying externally applied fields, such as deep brain stimulation (DBS) [3, 4]. The DBS technique has achieved substantial clinical success in treating movement disorders such as Parkinson’s disease, dystonia, and essential tremor, and is increasingly applied to epilepsy, chronic pain, obsessive–compulsive disorder, Tourette syndrome, depression, and other neuropsychiatric conditions, with emerging trials targeting circuits involved in Alzheimer’s disease, addiction, and post-traumatic stress disorder [5, 6]. Moreover, the EP is thought to play a role in the interaction of neighboring neurons that are not directly coupled through gap junctions or chemical synapses [7, 8, 9]. For instance, ephaptic interactions have been shown to contribute to inhibition [10] and synchronization [9] of Purkinje neurons, and to guide neocortical network activity [11]. In addition, modeling studies have demonstrated that endogenous fields perturb spike thresholds and timing and can entrain spiking [12, 13, 14].

In order to study these phenomena computationally, accurate and fully coupled models of the action potential (AP) and the associated EP is of considerable importance. Here, we apply a detailed computational model to quantify how external electrical stimulation modulates neuronal activity and to investigate how ephaptic coupling contributes to the interaction and synchronization of neighboring neurons.

Computational models of the EP surrounding neurons are most often based on a post-hoc, or open-loop, approach, in which the membrane potential and the EP are computed separately – and in that order [15, 16]. Solving the membrane equations under the assumption of a constant EP offers a substantial reduction in computational complexity. The EP can then be estimated from the transmembrane currents using analytical expressions based on line-source or point-source approximations [17, 18, 19]. These approximations provide useful estimates of local field potentials but neglect the bidirectional coupling between the cell membrane and the extracellular domain, which becomes important when cells are closely spaced, when strong external fields are applied, or when the aim is to understand ephaptic interactions.

Here, we follow the strategy introduced in [20] and further developed in [21, 22, 14], in which the dynamics of the extracellular (E) space, the cell membrane (M), and the intracellular (I) space are directly coupled through a system of partial differential equations – the EMI system. In addition to modeling the dynamics of neural tissue [20, 21, 22, 23, 14], the EMI system has also been applied to study the dynamics of small collections of cardiomyocytes [24, 25, 26].

In this study, we consider two neuron types: cerebellar Purkinje neurons and neocortical layer 5 pyramidal neurons. The membrane dynamics of these cells are described by models based on [27] and [28], respectively. The Purkinje neurons exhibit autonomous pacemaking activity, whereas the layer 5 pyramidal neurons remain quiescent in the absence of external input. Together, the two cell types span a broad range of electrophysiological properties, providing complementary cases for studying endogenous field interactions and externally driven responses.

In the presentation of the models, we show how the two neuron types respond to electrical stimulation applied to the cell membrane of the soma. The Purkinje neuron is subjected to a positive stimulation current, where increasing current strength reduces the number of APs per second. Conversely, the layer 5 pyramidal neuron is subjected to a negative stimulation current, which induces increasingly frequent APs as the current strength increases. Next, we study how the Purkinje and pyramidal neurons are modulated by an external current source as the distance to the source, the current amplitude, and the stimulation frequency are varied. In addition, we investigate ephaptic interactions between neighboring cells. First, we examine how rapidly two neighboring Purkinje neurons synchronize through ephaptic coupling as the cell distance is increased or as the extracellular conductivity is reduced. Moreover, we examine how the firing pattern of one layer 5 pyramidal neuron affects the firing pattern of a neigboring neuron through ephaptic coupling, and whether a neuron can directly trigger excitation in a neighboring neuron through ephaptic coupling.

Our aim is to contribute to the understanding of external stimulation of neurons, as used in DBS, and of ephaptic interactions in neural tissue, at the level of individual cells. Clearly, DBS acts on large populations of neurons [3, 4], but the basic mechanism is already present at the single-cell level. Similarly, neural interactions and synchronization through ephaptic coupling is typically studied in large neuronal populations, yet it remains important to understand the underlying mechanism between just two neurons [7, 8, 9, 13, 29]. This is the type of mechanistic understanding we aim to provide in the present study using the fully coupled EMI model.

## 2 Methods

### 2.1 The extracellular-membrane-intracellular (EMI) model

We represent the electrical potential of neurons and their surrounding extracellular space using the extracellular–membrane–intracellular (EMI) model [20, 21, 30]. The EMI model is given by the following system of equations:

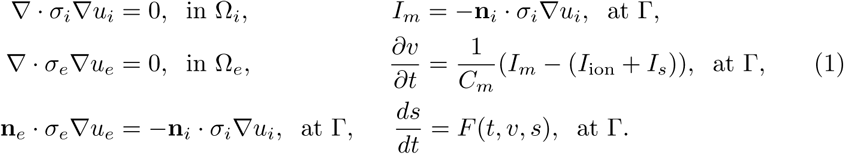

Here, *u_i_* and *u_e_* (in mV) are the intracellular and extracellular potentials, defined in the intracellular and extracellular domains, Ω*_i_* and Ω*_e_*, respectively. Moreover, *v* = *u_i_* − *u_e_* is the membrane potential given on the membrane, Γ, which is defined as the interface between Ω*_i_* and Ω*_e_*. In addition, *σ_i_* and *σ_e_* (in mS/cm) are the intracellular and extracellular conductivities, respectively, **n***_i_* and **n***_e_* are the outward pointing normal vectors of Ω*_i_* and Ω*_e_*, respectively, *C_m_* (in *µ*F/cm^2^) is the specific membrane capacitance, and *I_s_* (in *µ*A/cm^2^) is a stimulation current density. Unless otherwise stated, *I_s_* = 0. Time, *t*, is given in milliseconds (ms). Furthermore, *I*_ion_ (in *µ*A/cm^2^) is the current density through different types of ion channels on the cell membrane, *s* are gating variables and ionic concentrations involved in modeling these current densities, and *F* (*t, v, s*) model the dynamics of these additional state variables. For the model neurons considered in this study, the density of the different types of ion channels vary for different locations of the neuron, like the soma, the dendrites and the axon initial segment (AIS). This results in a spatially varying expression for *I*_ion_, which is described in more detail in the following subsections and in the Supplementary Information.

In addition, we consider axons partially covered by myelin. In the myelin covered part of the membrane, both *I*_ion_ and *C_m_* are set to zero, resulting in the no-flux boundary conditions

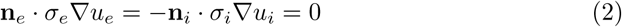

on the myelinated membrane.

In our simulations, we let the neurons be surrounded by approximately 500 *µ*m of extracellular space in spatial each direction. On the outer boundary of the extracellular domain, we apply the Dirichlet boundary condition

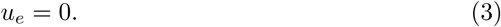

### 2.2 Cerebellar Purkinje neuron model

We model cerebellar Purkinje neurons using the EMI modeling setup described in [14], based on the cable model formulation from [27]. The Purkinje neuron is divided into seven different morphological parts, the dendrites, the soma, the axon initial segment (AIS), a ParaAIS, three Ranvier nodes separated by myelinated membrane sections and an axon collateral (see Figure 1A). The ion channel densities for each of these sections are described in the Supplementary Information. The three-dimensional (3D) finite element mesh of a neuron and the surrounding extracellular space is illustrated in Figure 1B.

**Figure 1:**
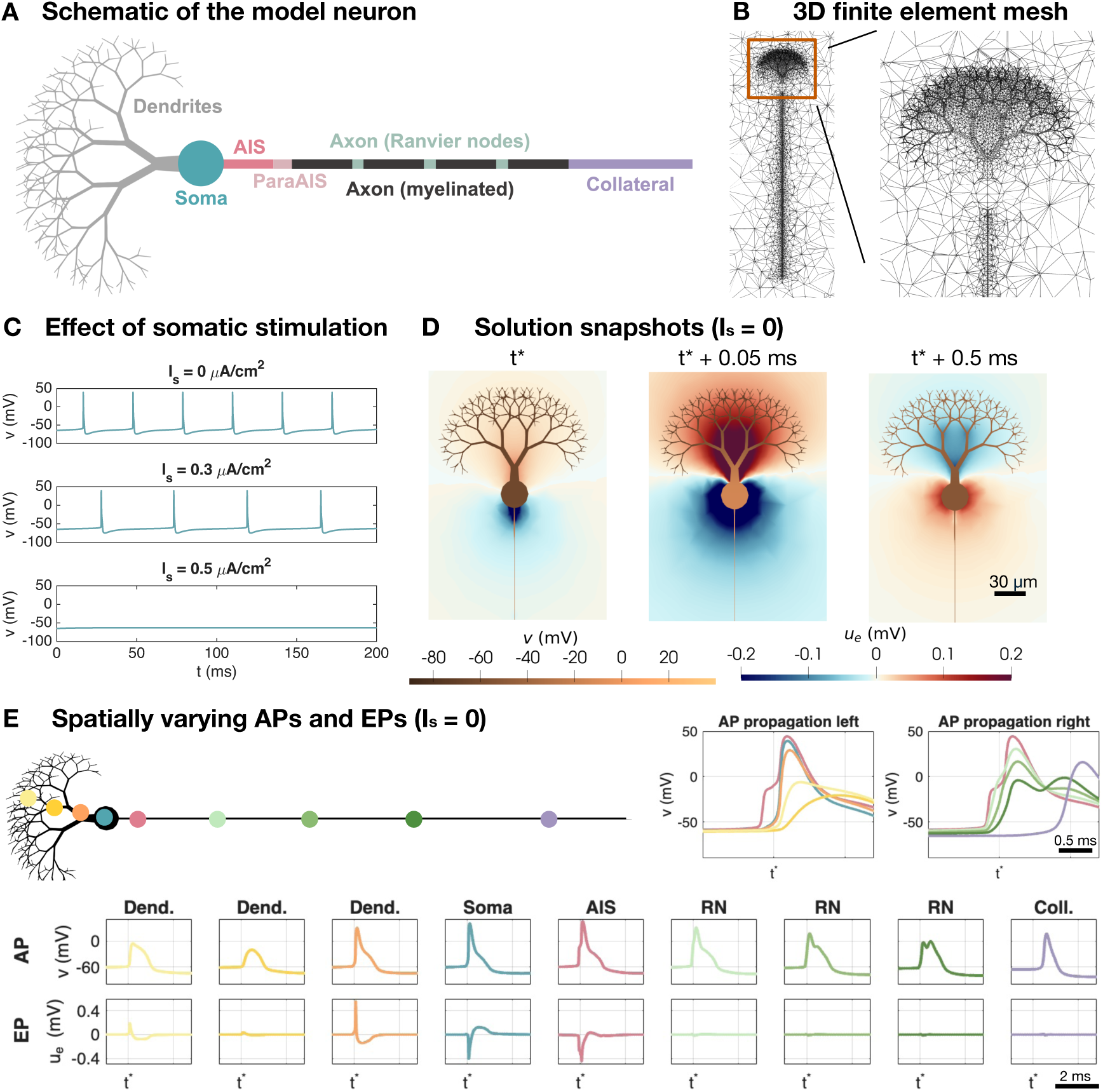
Setup of the cerebellar Purkinje neuron model. A: Schematic of the different geometrical parts of the model neuron. The membrane model differs between these parts as described in the Supplementary Information. B: Illustration of the 3D finite element mesh used in the EMI model simulations. C: Effect of somatic stimulation (*I_s_*) on the somatic membrane potential. For *I_s_* = 0, the cell fires APs spontaneously. For *I_s_* = 0.3 *µ*A/cm^2^, the firing frequency is reduced and for *I_s_* = 0.5 *µ*A/cm^2^, the AP firing ceases. D: Solution snapshots of the membrane potential and EP in the case of no somatic stimulation. E: APs and EPs at different locations of the neuron. The AP is initiated in the AIS and propagates to the left through the soma and dendrites and to the right through the axon.

In the absence of stimulation, the Purkinje neuron fires spontaneous APs. These are initiated in the AIS and propagate left through the soma and dendrites and right through the axon (see Figure 1E). In the soma and AIS clear negative EPs are generated during AP firing, whereas positive EPs are generated near the dendrites. Figure 1D provide spatial solution snapshots of the membrane potential and the surrounding EP at three points in time during AP firing.

If a positive stimulation current, *I_s_*, is applied in the soma of the Purkinje neuron model, the frequency of the AP firing is reduced, and if the stimulation is sufficiently strong (e.g., *I_s_* = 0.5 *µ*A/cm^2^), the AP firing is completely suppressed (see Figure 1C).

### 2.3 Neocortical layer 5 pyramidal neuron model

The model for neocortical layer 5 pyramidal neurons is set up in a similar manner as the model for cerebellar Purkinje neurons (see Figure 2). The model for the membrane dynamics is based on [28], and the specific parameter values are specified in the Supplementary Information.

**Figure 2:**
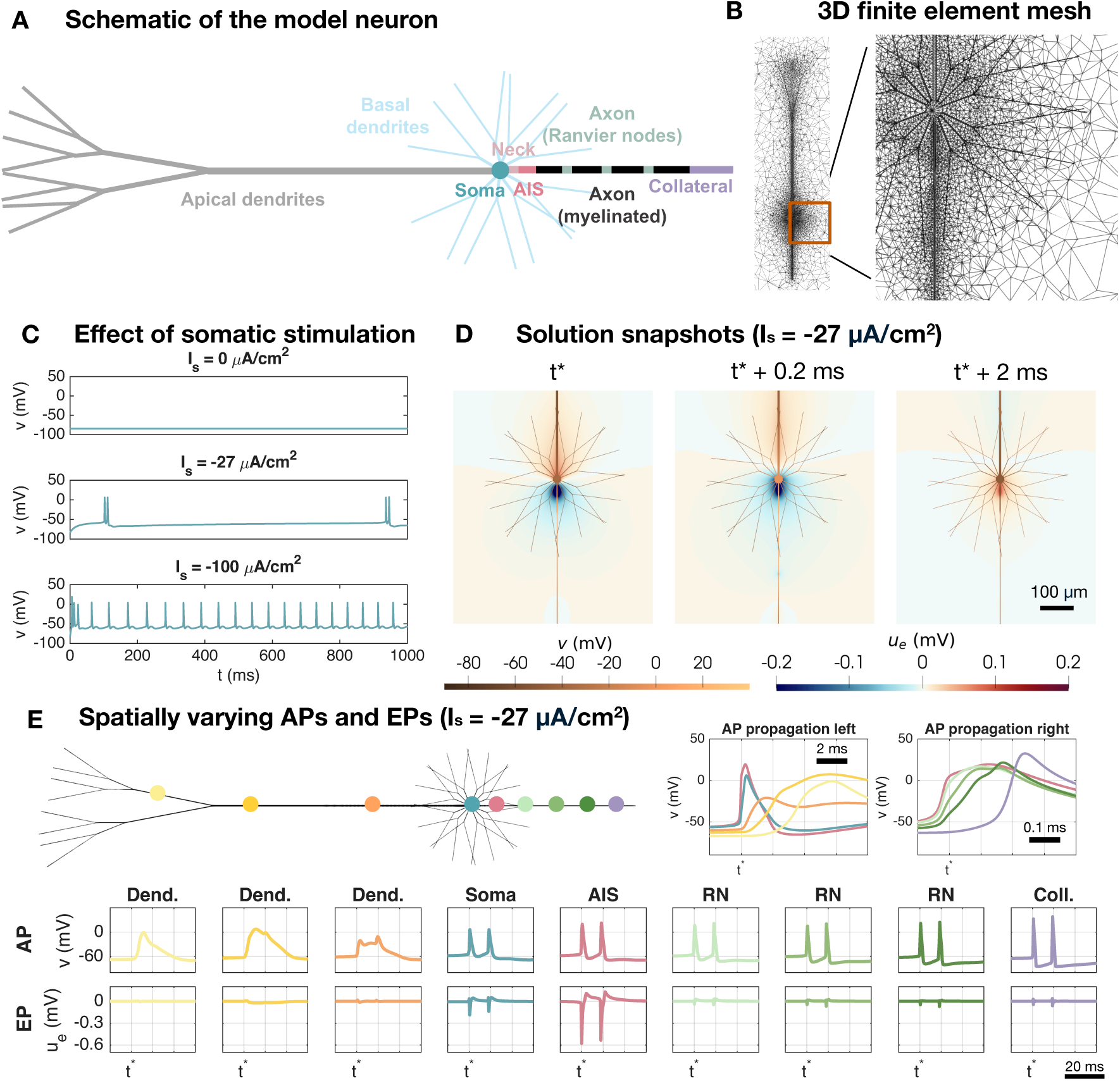
Setup of the neocortical layer 5 pyramidal neuron model. A: Schematic of the different geometrical parts of the model neuron. The membrane model differs between these parts as described in the Supplementary Information. B: Illustration of the 3D finite element mesh used in the EMI model simulations. C: Effect of somatic stimulation (*I_s_*) on the somatic membrane potential. For *I_s_* = 0, the cell do not fire APs, but for negative somatic stimulation, APs are fired. D: Solution snapshots of the membrane potential and EP in the case of *I_s_* = *−*27 *µ*A/cm^2^. E: APs and EPs at different locations of the neuron in the case of *I_s_* = *−*27 *µ*A/cm^2^. The AP is initiated in the AIS and propagates to the left through the soma and dendrites and to the right through the axon.

The pyramidal neuron morphology consists of eight different parts, the apical dendrites, the basal dendrites, the soma, the axon neck between the soma and the AIS, the AIS, three Ranvier nodes separated by myelinated membrane sections and an axon collateral (see Figure 2A). The channel densities and geometry for each of these different sections are specified in the Supplementary Information. In Figure 2B, the 3D finite element mesh used to represent a pyramidal neuron and its surrounding extracellular space is illustrated.

In the absence of stimulation, the pyramidal neuron do not fire APs, but when the soma is stimulated by a sufficiently strong negative stimulation current, *I_s_*, APs are fired (see Figure 2C). When the strength of *I_s_* is further increased, the frequency of the AP firing increases.

In response to a somatic stimulation (*I_s_* = −27 *µ*A/cm^2^), the AP is first initiated in the AIS and propagates left through the axon neck, soma and dendrites, and right through the axon, as depicted in Figure 2E. In the apical dendrite, the AP lasts considerably longer than in the soma and axon. More specifically, two AP spikes occur in the soma and axon during one spike in the apical dendrite for this somatic stimulation strength. During AP firing there are clear negative EP spikes near the soma and AIS, and the largest spikes are generated near the AIS, consistent with experimental observations in [31]. Figure 2D shows spatial solution snapshots of the membrane potential of the neuron and the surrounding EP near the soma and AIS during AP firing.

### 2.4 Stimulation protocols

In our simulations, we consider three different types of stimulation.

**I: Constant somatic stimulation** The first considered type of stimulation is a constant membrane current density, *I_s_* (in *µ*A/cm^2^), applied to the somatic part of the membrane. This membrane current density is included in the EMI model as described in (1). A positive *I_s_* reduces the neuron’s excitability, whereas a negative *I_s_* increases the excitability of the neuron (see Figures 1C and 2C).

**II: Constant extracellular stimulation** A second type of considered stimulation is an extracellular point source, representing extracellular stimulation through an electrode. This point source is represented by a sphere cut out of the mesh of the extracellular domain. On the surface of this sphere, a Neumann boundary condition of the form

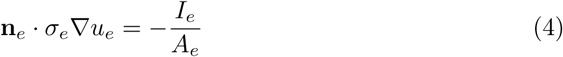

is applied, where *I_e_* (in *µ*A) is the stimulation strength, *A_e_* (in cm^2^) is the surface area of the sphere, and **n***_e_* is the outward pointing normal vector of the extracellular domain (i.e., pointing towards the center of the stimulation sphere). For a positive *I_e_*, a negative EP is emanating from the stimulation sphere. The stimulation sphere is located in the vicinity of the AIS and the distance will be reported for each simulation. The EP surrounding the Purkinje neuron at rest as a result of a constant extracellular stimulation is illustrated for a few choices of *I_e_* in Figure 3A. In addition, the EP at an AIS membrane point as a function of the electrode-to-AIS distance is provided in Figure 3B in a similar manner as in Figure 1B in [29] and Figure 1C in [7].

**Figure 3:**
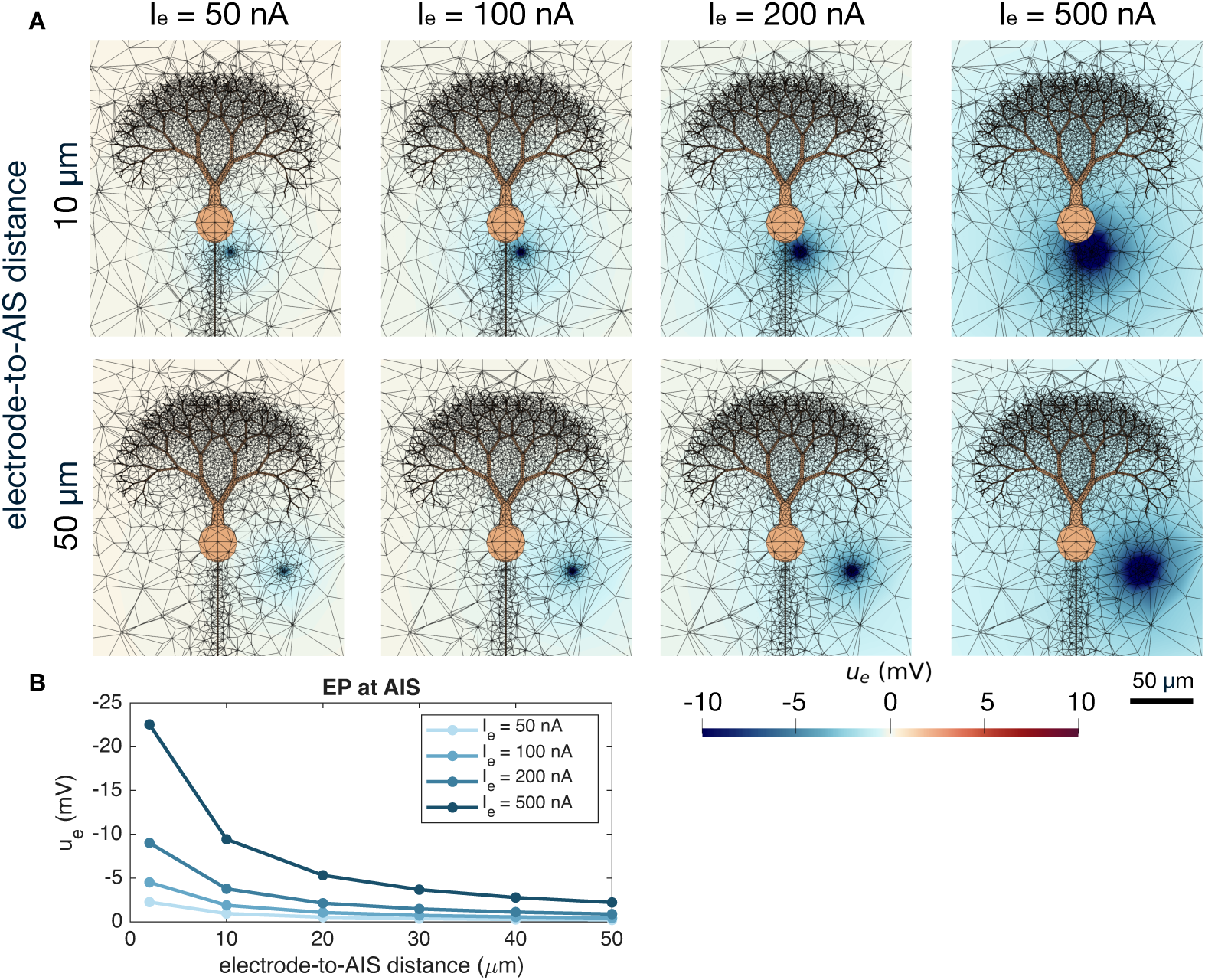
Extracellular potential generated by a constant extracellular stimulation source near a cerebellar Purkinje neuron. A: Extracellular potential in a sheet in the center of the domain in the *z*-direction. In addition, the finite element mesh is indicated by black lines and the neuron is shown in orange. In the upper panel, the distance between the stimulation source and the AIS is 10 *µ*m, and in the lower panel the distance is 50 *µ*m. B: Extracellular potential measured at an AIS membrane point of the Purkinje neuron as a function of the distance to the electrode.

**III: Sinusoidal extracellular stimulation** The third considered stimulation type is a sinusoidal extracellular stimulation changing strength and sign with time during the simulation. This stimulation is implemented in the same manner as the constant extracellular stimulation, except that the boundary condition on the stimulation sphere is given by

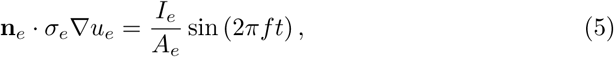

where *I_e_*(in *µ*A) is the stimulation strength, *A_e_*(in cm^2^) is the surface area of the stimulation sphere, *f* (in Hz) is the stimulation frequency, and *t* (in seconds) is time.

### 2.5 Numerical methods

We solve the EMI system of equations numerically using the spatial and temporal operator splitting technique introduced in [32] with a global time step of Δ*t* = 0.01 ms. We apply the MFEM finite element library [33, 34] with first-order elements and a mesh generated using Gmsh [35]. For the non-linear membrane model dynamics, we use the first order Rush Larsen method [36] with code generated using the Gotran software [37] and a time step of Δ*t* = 0.001 ms for the Purkinje neuron and Δ*t* = 0.01 ms for the pyramidal neuron.

## 3 Results

In this section, we report results of our EMI model simulations of extracellular stimulation and ephaptic interactions of neurons. We begin by considering a single neuron in the vicinity of an extracellular stimulation source. We consider either a single cerebellar Purkinje neuron or a single neocortical layer 5 pyramidal neuron. The purpose of these simulations is to investigate how alterations in the EP (caused by, e.g., extra-cellular stimulation or by other firing neurons) can affect single-neuron dynamics. We start by considering subthreshold effects on the neuronal membrane potential. Then we move on to investigate how extracellular stimulation can modulate the number of AP spikes and the timing of AP spikes. Next, we focus on two neigbouring neurons that are not connected by synapses or gap junctions and investigate ephaptic interactions between the two cells.

### 3.1 Subthreshold membrane potential effects induced by weak extracellular stimulation

As a first investigation into the effects of extracellular stimulation on neurons, we consider a non-firing cerebellar Purkinje neuron and a non-firing neocortical layer 5 pyramidal neuron located near a sinusoidal extracellular stimulation source with relatively weak stimulation amplitudes. The Purkinje neuron is made quiescent by a somatic stimulation current of *I_s_* = 1 *µ*A/cm^2^, while no somatic stimulation is applied for the pyramidal neuron. The extracellular stimulation source is located 20 *µ*m from the AIS of the neurons. Figure 4 shows the membrane potential and the EP in an AIS membrane point of the two neurons. We observe that the extracellular stimulation yields an EP of up to about ± 0.5 mV and ± 1 mV for the stimulation amplitudes of *I_e_* = 50 nA and *I_e_* = 100 nA, respectively, at the AIS membrane. This results in a similarly sized alteration in the AIS membrane potential of the two cells. These subthreshold membrane potential fluctuations are present for different applied stimulation frequencies (1 Hz, 5 Hz, and 30 Hz), consistent with previous experimental findings [7, 29].

**Figure 4:**
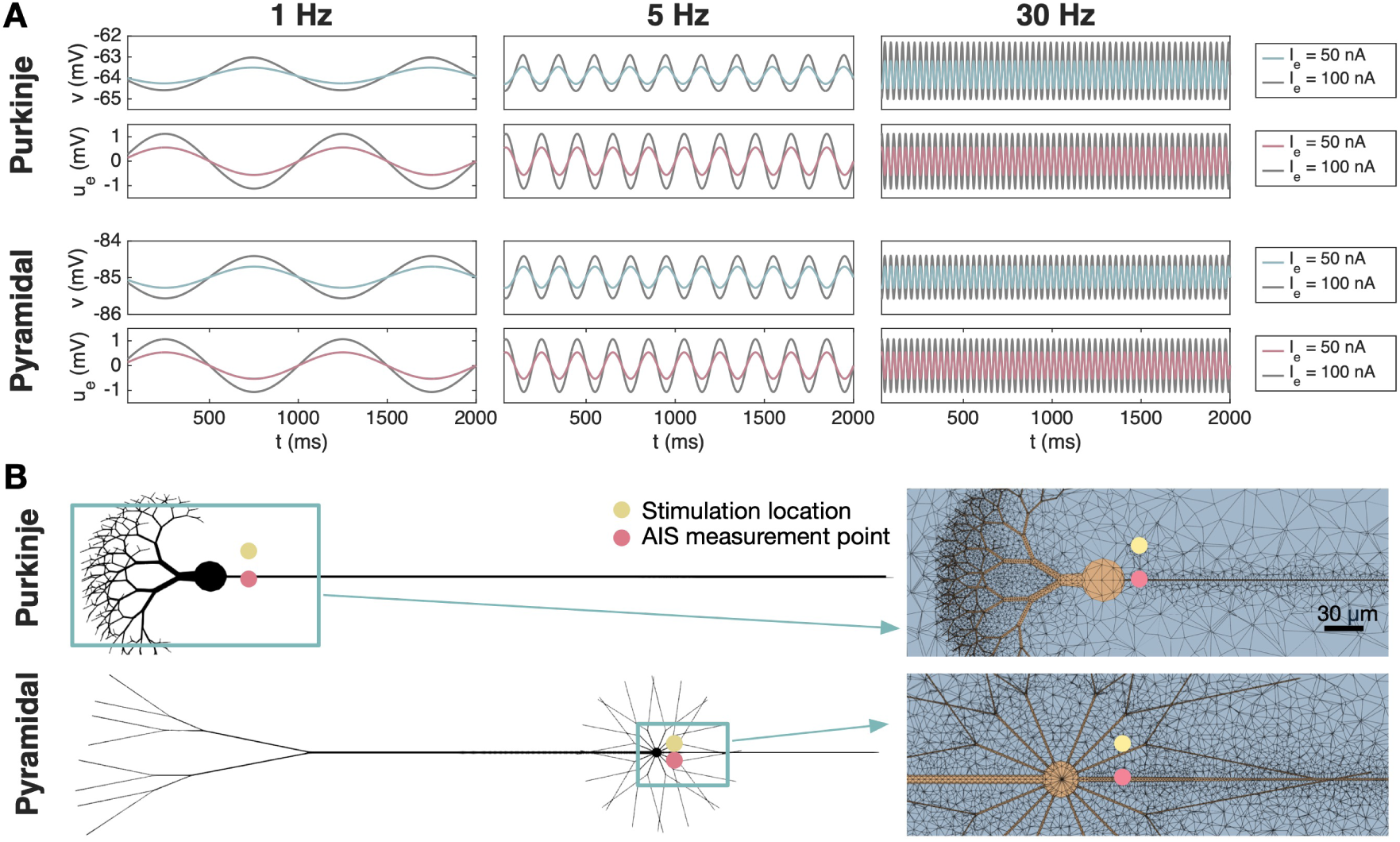
Subthreshold effects at the AIS of a cerebellar Purkinje neuron and a neocortical layer 5 pyramidal neuron under sinusoidal extracellular stimulation. For the pyramidal neuron, no somatic stimulation (*I_s_*) is applied, whereas for the Purkinje neuron, *I_s_* = 1 *µ*A/cm^2^ is used to maintain a stable resting membrane potential. A: Membrane potential, *v*, and extracellular potential, *u_e_*, shown at an AIS membrane point for three different stimulation frequencies (1 Hz, 5 Hz, 30 Hz) and two different stimulation strengths (*I_e_* = 50 nA and *I_e_* = 100 nA). B: Illustration of the simulation setup and a part of the 3D mesh used in the simulations. The AIS measurement point is indicated by a pink circle and the stimulation electrode is illustrated by a yellow circle. The distance between the stimulation electrode and the AIS is 20 *µ*m.

### 3.2 Extracellular stimulation modulates neuronal spikes

Next, we investigate how extracellular stimulation modulates the dynamics of firing neurons.

#### 3.2.1 Constant extracellular stimulation modulates the number of AP spikes

In Figure 5, we examine how a constant extracellular stimulation affects the number of AP spikes during a 500 ms simulation. Moreover, we investigate how this number is influenced by the extracellular stimulation strength, *I_e_*, and the distance between the stimulation source and the neuron AIS.

**Figure 5:**
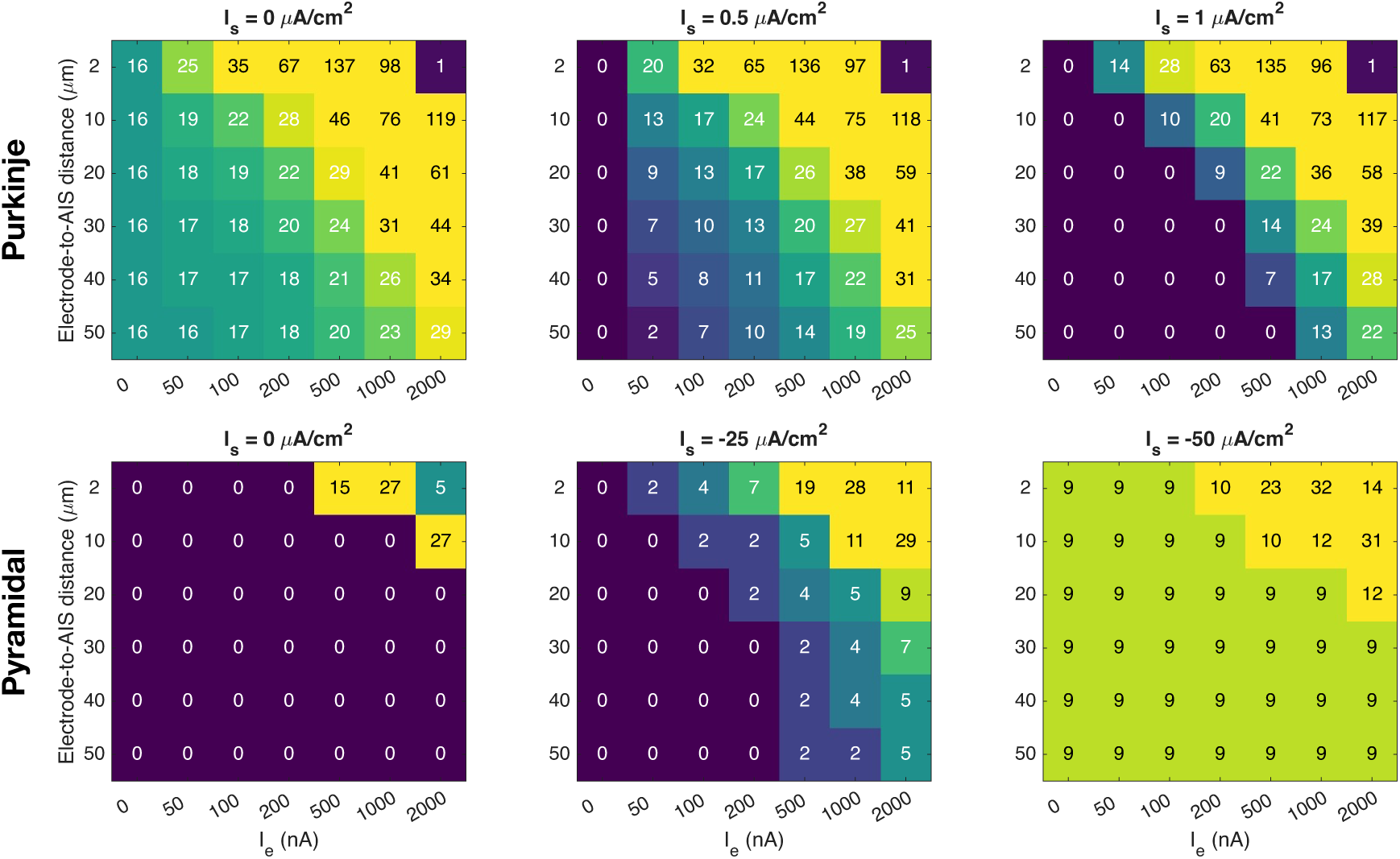
Effect of constant extracellular stimulation on the number of spikes for a cerebellar Purkinje neuron and a neocortical layer 5 pyramidal neuron. For each extracellular stimulation strength, *I_e_*, and electrode-to-AIS distance, the number of somatic spikes during a 500 ms simulation is reported. Three different levels of somatic stimulation, *I_s_*, are considered for each cell type.

In the upper panel of Figure 5, extracellular stimulation is applied near a Purkinje neuron, and in the left panel, no somatic stimulation is applied. If no extracellular stimulation is applied, the neuron fires 16 spikes during the 500 ms simulation. However, when a sufficiently strong extracellular stimulation is applied sufficiently close to the neuron, the spike frequency is increased. The number of spikes increases further as the extracellular stimulation strength is increased and as the distance between the stimulation source and the neuron decreases. Yet, when a very strong stimulation is applied close to the neuron (upper right corner), the stimulation is so strong that the neuron fails to repolarize, so there is no rapid AP firing. Moreover, for a distance of 2 *µ*m and *I_e_* = 1000 nA, the neuron approaches the no-repolarization state causing a reduced number of spikes compared to the case of *I_e_* = 500 nA.

In the middle and right upper panels of Figure 5, a positive stimulation current, *I_s_*, is included across the membrane of the soma, making the cell quiescent in the absence of extracellular stimulation. However, if a sufficiently strong extracellular stimulation is applied sufficiently near the neuron, AP firing is initiated and, like in the left panel, the firing frequency increases as the extracellular stimulation strength is increased or the distance between the stimulation source and the neuron AIS is reduced. We also observe that, as expected, a stronger extracellular stimulation is required to initiate AP firing when the positive somatic stimulation is increased because this makes the cell less excitable (compare middle and right upper panels of Figure 5).

In the lower panel of Figure 5, we consider a layer 5 pyramidal neuron near the stimulation source. In the absence of extracellular stimulation, this neuron does not fire APs if there is no somatic stimulation or if a relatively weak somatic stimulation of *I_s_* = −25 *µ*A/cm^2^ is applied. Yet, AP firing may be initiated if a sufficiently strong extracellular stimulation is applied sufficiently near the neuron (see lower left and middle panels of Figure 5). As expected, AP firing is more easily initiated if some negative somatic stimulation is present (middle panel) compared to the case of no somatic stimulation (left panel). In the right panel, the somatic stimulation is strong enough to initiate AP firing in the absence of extracellular stimulation, but the cell fires more APs if a sufficiently strong extracellular stimulation is applied sufficiently near the neuron AIS. Nevertheless, the layer 5 pyramidal neuron generally appears to require a stronger extracellular stimulation to affect AP firing than what is required to modulate the firing of the Purkinje neuron.

#### 3.2.2 Sinusoidal extracellular stimulation modulates the timing of AP spikes

After considering a constant extracellular stimulation source, we now focus on AP spikes modulated by extracellular stimulation of a sinusoidal shape. Figure 6A shows the membrane potential and EP of a cerebellar Purkinje neuron near an extracellular stimulation source, and Figure 6B shows the simulation setup. We consider four different stimulation frequencies (1 Hz, 5 Hz, 30 Hz and 100 Hz) and three different stimulation strengths (*I_e_* = 100 nA, *I_e_* = 200 nA, and *I_e_* = 1000 nA) applied 20 *µ*m from the AIS. No somatic stimulation is applied. In the left panel of Figure 6A, a 1 Hz stimulation frequency is considered. We observe that this results in an EP of sinusoidal shape at the AIS with small extracellular spikes during AP firing (see the boxed inset in the upper left panel). For a relatively weak stimulation (100 nA), the AP spikes appear to occur more frequently when the EP is negative and less frequently when the EP is positive. When the amplitude of the extracellular stimulation is increased this tendency is even more pronounced, with no AP firing during the most positive values of the EP. For 5 Hz stimulation, similar behavior is observed. The 30 Hz stimulation, on the other hand, is more similar to the inherent firing frequency of the Purkinje neuron, and the cell appears to simply fire once each time the EP is negative. For the 100 Hz frequency, the stimulation effect seem to be limited for the low stimulation amplitude, whereas for the higher amplitudes the cell fires every (*I_e_* = 1000 nA) or every other (*I_e_* = 200 nA) time the EP is in a negative phase.

**Figure 6:**
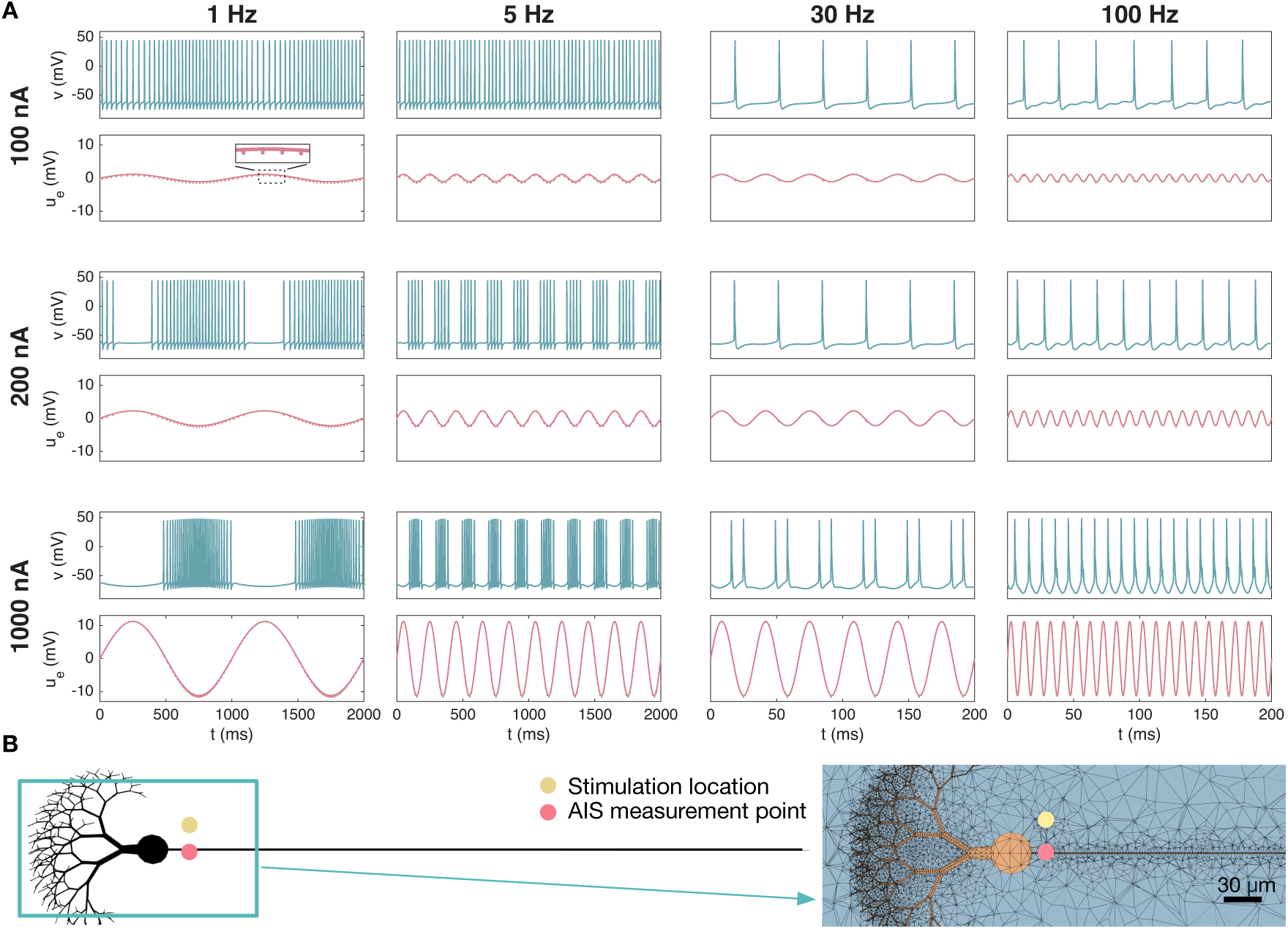
Effect of sinusoidal extracellular stimulation on the spike timing of a cerebellar Purkinje neuron. A: The membrane potential, *v*, and the extracellular potential, *u_e_*, at an AIS membrane point for four different stimulation frequencies and three different stimulation strengths. In the extracellular potential traces, we observe small perturbations caused by the AP firing, which are magnified in the boxed inset for the 1 Hz, 100 nA case. Note that the scaling of the *x*-axis is different between the two cases of high frequency and the two cases of low frequency. B: Illustration of the simulation setup and a part of the 3D mesh used in the simulations. The distance between the stimulation electrode and the AIS is 20 *µ*m and no somatic stimulation, *I_s_*, is applied.

Figure 7 shows the results of similar simulations of a layer 5 pyramidal neuron in the vicinity of extracellular stimulation. In this case, we apply a somatic stimulation current of −27 *µ*A/cm^2^, which is strong enough to trigger AP firing of the neuron, but only by a narrow margin. In this case, we observe that the frequency and strength of the extracellular stimulation has an effect on the timing and number of AP spikes, but the connection between extracellular stimulation and AP firing appears a bit less straightforward than for the Purkinje neuron.

**Figure 7:**
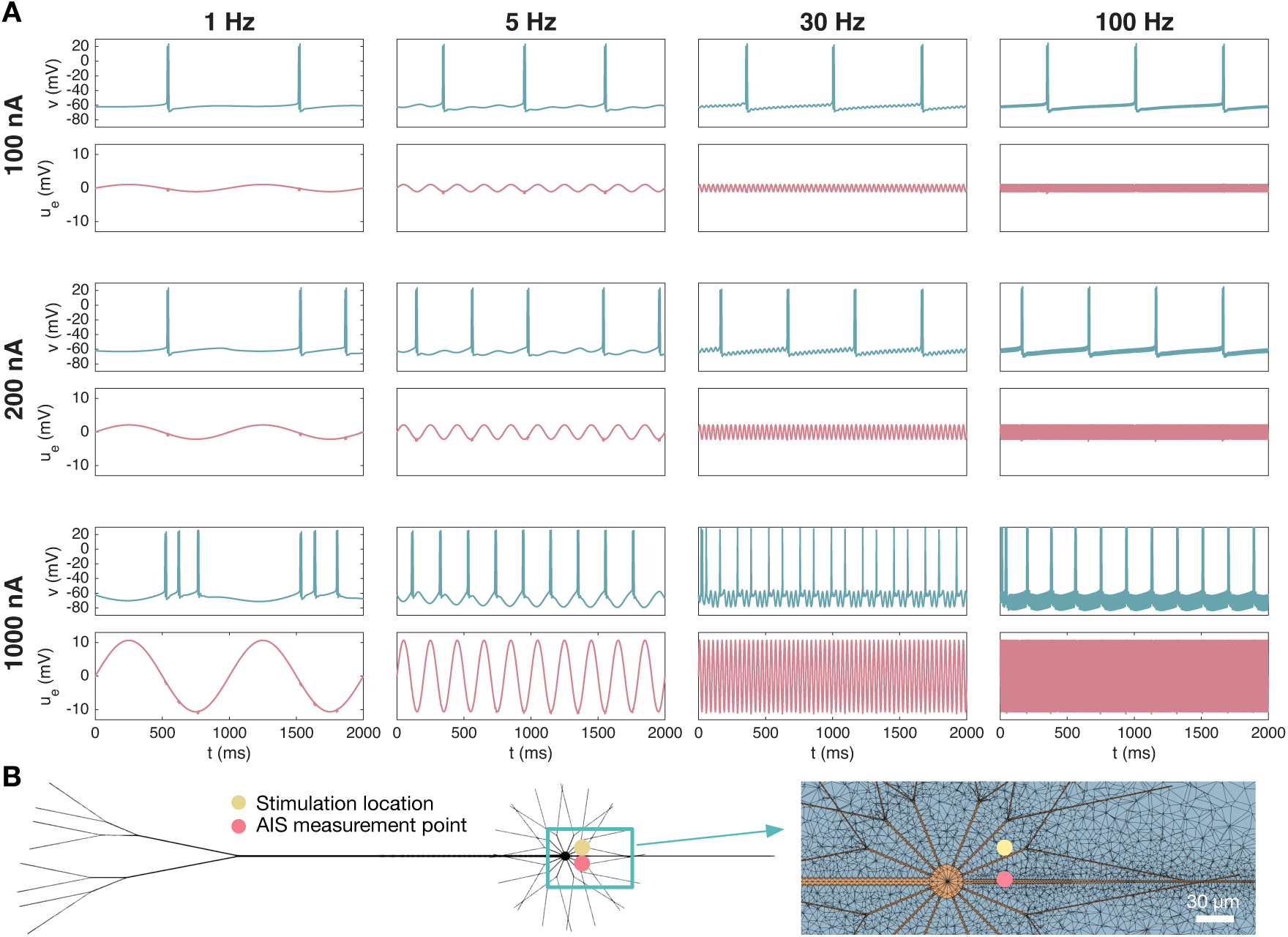
Effect of sinusoidal extracellular stimulation on the spike timing of a neocortical layer 5 pyramidal neuron. A: The membrane potential, *v*, and the extracellular potential, *u_e_*, at an AIS membrane point for four different stimulation frequencies and three different stimulation strengths. B: Illustration of the simulation setup and a part of the 3D mesh used in the simulations. The distance between the stimulation electrode and the AIS is 20 *µ*m and, and the somatic stimulation, *I_s_*, is set to *−*27 *µ*A/cm^2^.

### 3.3 Synchronization of two cerebellar Purkinje neurons

The next stage in our investigations is the study of ephaptic effects between two neurons. As a first example of such effects, we consider ephaptic synchronization of two firing cerebellar Purkinje neurons following up our preliminary analysis of this topic in [14].

#### 3.3.1 Synchronization of spiking Purkinje neurons depends on cell distance

In Figure 8, we explore potential synchronization of two firing Purkinje neurons for three different cell distances. From left to right, the minimal distance between the AIS of the two neurons is 10 *µ*m, 5 *µ*m and 1 *µ*m. The axon of the neurons are shaped to bend towards each other in order to achieve these relatively small cell distances (see Figure 8A). No somatic or extracellular stimulation is applied, and the cells beat spontaneously. In order to study synchronization, we manipulate the initial part of the simulation such that the left neuron (denoted as Cell 1) fires APs approximately 5 ms before the right neuron (denoted as Cell 2). The cells are not connected by gap junctions or synapses, but the EP generated when one cell fires subtly alters the membrane potential of the other cell. In Figure 8, we observe that these subtle effects eventually cause the two neurons to start firing APs in synchrony if they are located sufficiently close. This was also observed in [14].

**Figure 8:**
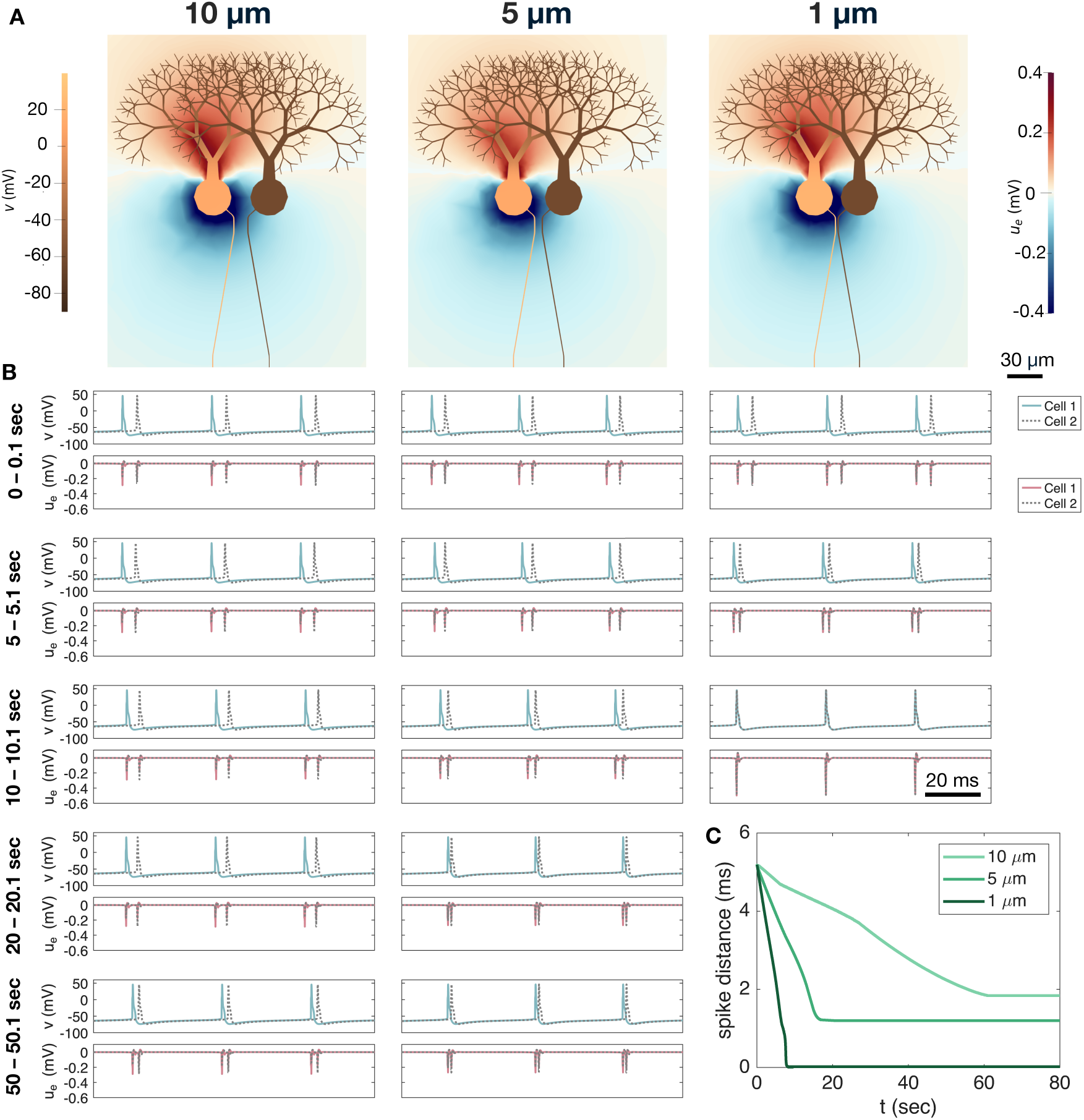
Synchronization of two cerebellar Purkinje neurons for different cell distances. The shortest distance between the AISs of the two neurons is 10 *µ*m, 5 *µ*m and 1 *µ*m in the left, middle, and right panels, respectively. A: Solution snapshots of the extracellular potential (*u_e_*) and the membrane potential (*v*, color shown inside neuron) during AP firing of the left cell (Cell 1). B: Membrane potential and extracellular potential in an AIS membrane point of each cell. Each panel row focuses on a different time interval during the simulation, and the columns correspond to the same three cell distances as in Panel A. Note that for a cell distance of 1 *µ*m, the cells are completely synchronized after 10 seconds. Therefore, the plots for last time interval is omitted for this cell distance. C: Temporal distance between AP spikes for the two cells as a function of time for the three different cell distances. Synchronization is considerably faster and the synchronized spike difference is considerably smaller for a short cell distance.

Figure 8A shows snapshots of the EP and membrane potential solutions during AP firing of Cell 1 for the three different cell distances. We observe that a negative EP emanates from the AIS and soma of Cell 1, causing a negative EP at the membrane of Cell 2. Note that during AP firing of Cell 2 a similar EP is generated from that cell. Figure 8B shows the membrane potential and EP at an AIS membrane point of each cell. The panel columns correspond to the three different considered cell distances and the rows represent different time intervals during a long simulation. In the upper row, we consider the solutions at the start of the simulation. In this case, Cell 2 fires APs approximately 5 ms after Cell 1. In the next row, the cells have been firing APs next to each other for five seconds. For a 10 *µ*m cell distance, the solutions appear quite similar as those recorded in the start of the simulation, but for the smaller cell distances, the distance between the AP spikes of the two cells have clearly been reduced. In the third row, the cells have been firing APs next to each other for ten seconds. Now, there is no longer any visible distance between the AP spikes for a 1 *µ*m cell distance, implying that the cells are now synchronized. Note also that the amplitude of the EP spikes approximately doubles in this case because the EPs generated by the two cells add up. After twenty seconds of simulation, the distance between spikes is about 1 ms for the 5 *µ*m cell distance. However, after fifty seconds, the spike distance appears to be approximately the same, indicating that the cells have reached a stable state of synchrony.

In Figure 8C, the results of Panel B are summarized. Here, the temporal distance between the spikes of the two cells are plotted as a function of time for the three considered cell distances. We observe that once the cells have reached a stable state of synchrony, the distance between spikes do not start to increase again; the cells remain synchronized. In addition, synchronization occurs considerably faster as the cell distance is reduced and the stable distance between spikes is shorter as the cell distance is reduced. This is consistent with the experimental observation of improved cerebellar Purkinje neuron synchronization for reduced AIS distances in [9].

#### 3.3.2 Rapid synchronization of Purkinje neurons in presence of high extracellular resistivity

In Figure 9, we fix the AIS distance between the two cells at 1 *µ*m and consider the case of increased extracellular resistivity. Since resistivity is the reciprocal of conductivity, this corresponds to a reduced extracellular conductivity, *σ_e_*. We consider the default value of *σ_e_* = 3 mS/cm [38] as well as three levels of reduction: *σ_e_/*2, *σ_e_/*5, and *σ_e_/*10. Like in Figure 8, the initial part of the simulation is manipulated such that Cell 1 (left neuron) initially fires APs approximately 5 ms before Cell 2 (right neuron).

**Figure 9:**
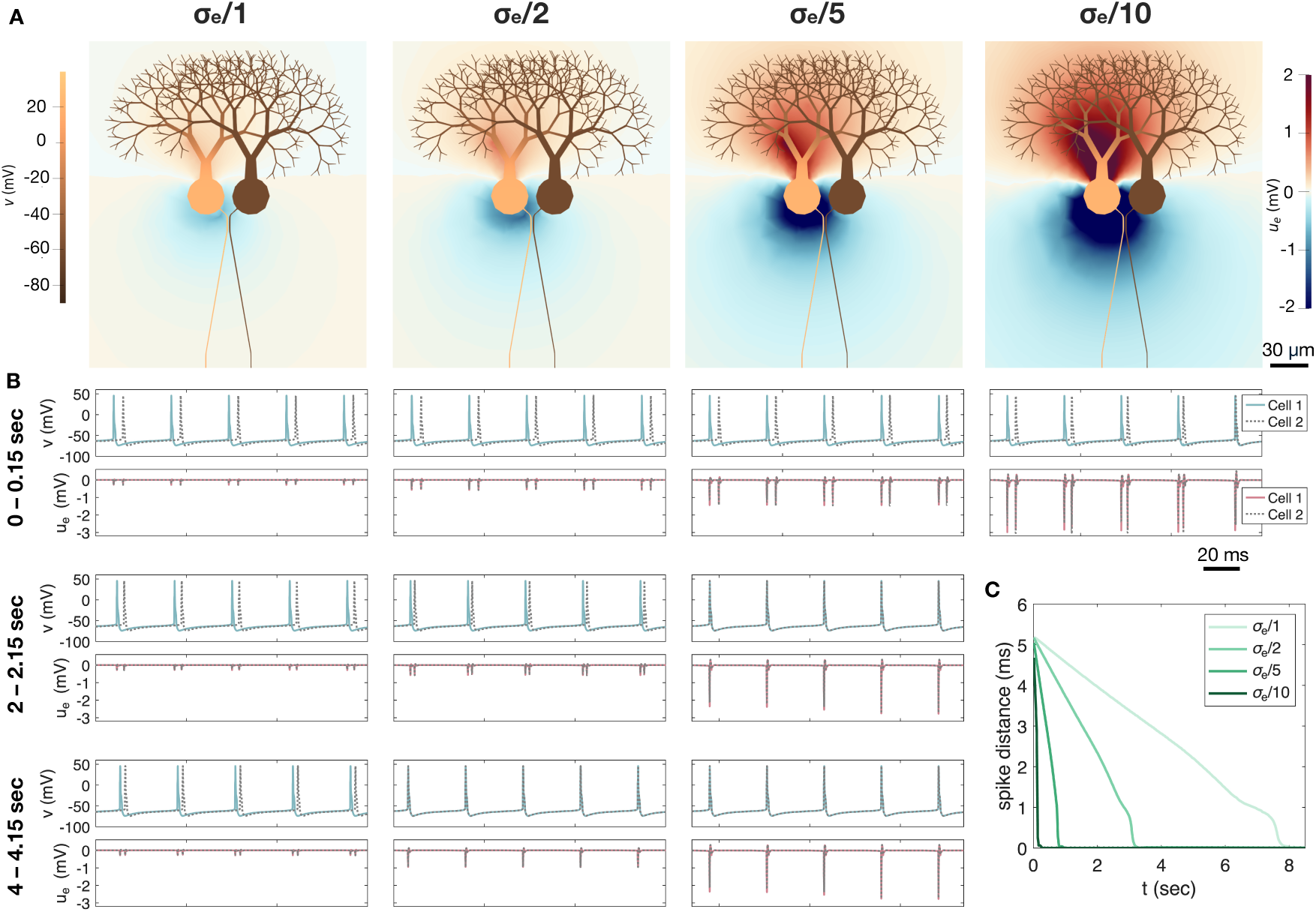
Synchronization of two cerebellar Purkinje neurons in the presence of reduced extracellular conductivity, *σ_e_*. The AISs of the two neurons are separated by a minimum of 1 *µ*m of extracellular space. A: Solution snapshots of the extracellular potential (*u_e_*) and the membrane potential (*v*) during AP firing of the left cell (Cell 1) for four different values of *σ_e_*. The default value of *σ_e_* = 3 mS/cm is divided by 1, 2, 5, or 10. B: Membrane potential and extracellular potential in an AIS membrane point of each cell at different time intervals during the simulation. We consider the same four values of *σ_e_* as in Panel A. Note that for *σ_e_/*10, the cells are almost completely synchronized after only five spikes. Therefore, the plots for later time intervals are omitted for this value of *σ_e_*. C: Temporal distance between AP spikes for the two cells as a function of time for the four different values of *σ_e_*. Synchronization is considerably faster for reduced values of *σ_e_*.

Figure 9A shows snapshots of the membrane potential and EP during Cell 1 firing. We observe that the magnitude of the EP is considerably larger for a reduced extra-cellular conductivity. Note again, that a similar EP emerge from Cell 2 when that cell fires APs. Figure 9B shows the membrane potential and EP in an AIS membrane point of the two cells at a few different time intervals. When *σ_e_* is reduced by a factor of 10, we observe that the two cells are almost completely synchronized already after only five AP spikes. For the remaining values of *σ_e_*, synchronization takes a bit longer, but it occurs faster for small values of *σ_e_*. In Figure 9C, these results are summarized by plotting the distance between spikes for the two cells as a function of time. We observe that for the default value of *σ_e_*, synchronization takes about 8 seconds, for *σ_e_/*2, it takes a little more than 3 seconds, and for *σ_e_/*5 it takes less than a second.

### 3.4 Ephaptic interactions between two neocortical layer 5 pyramidal neurons modulate spike timing

We now consider the case of ephaptic interactions between two neocortical layer 5 pyramidal neurons. In this case, we examine how the AP firing of a cell receiving substantial somatic stimulation modulates the AP firing of a nearby cell receiving weak somatic stimulation. Figure 10 shows such ephaptic interactions between two spiking pyramidal neurons. Like for the Purkinje neurons, we let the axon of the pyramidal neurons bend towards each other such that the minimum distance between the AIS of the two cells is 1 *µ*m (see Figure 10A). The two cells are not connected by gap junctions or synapses. In Figure 10A, we show solution snapshots for a few time points during AP firing of the left cell (Cell 1). First, a negative EP is generated close to the AIS, and later towards the soma. Then, after a few milliseconds, a positive EP is present close to the soma and the AIS.

**Figure 10:**
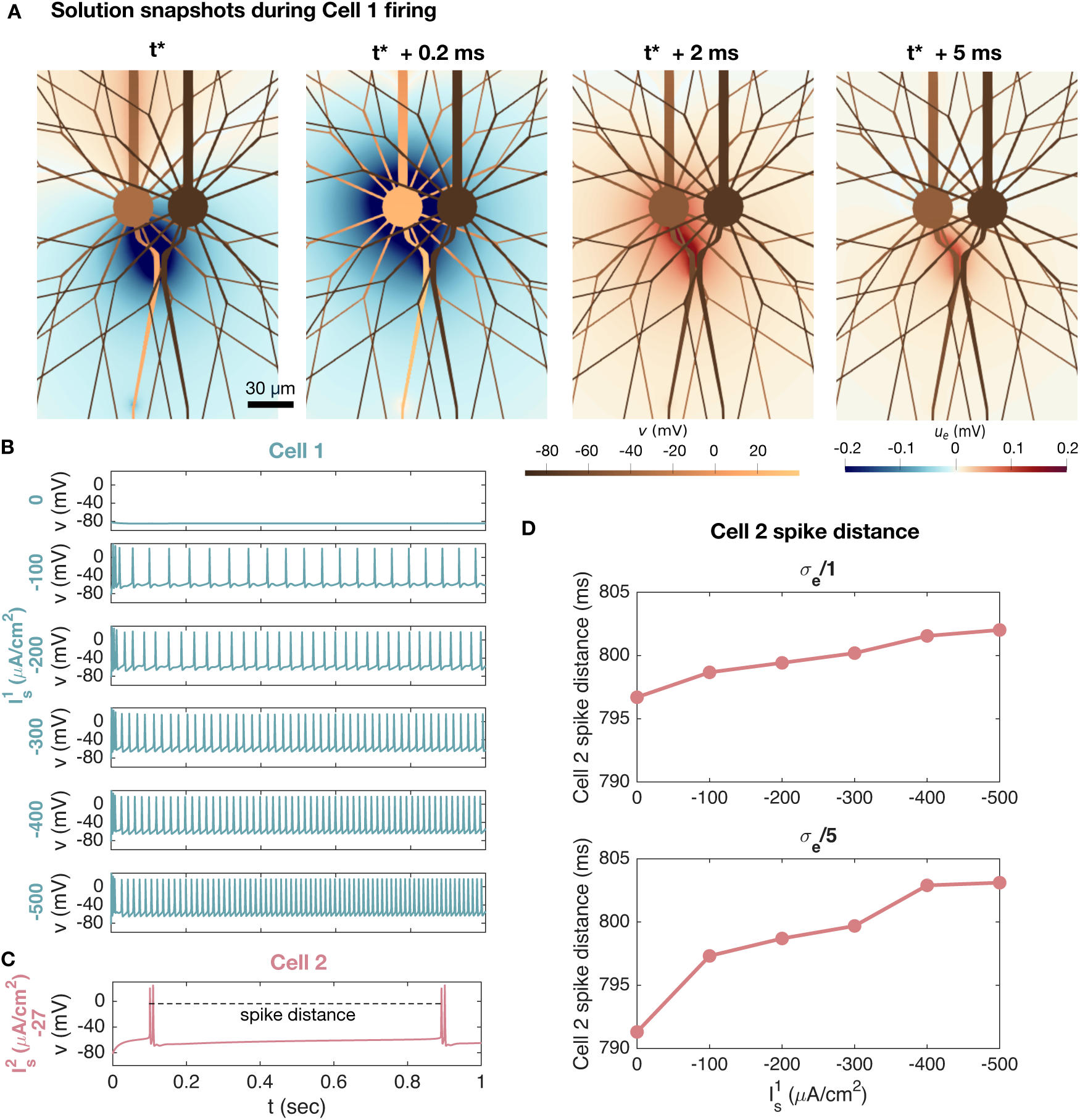
Ephaptic interactions between two neocortical layer 5 pyramidal neurons. The AISs of the two neurons are separated by 1 *µ*m of extracellular space. A: Solution snapshots of the extracellular potential (*u_e_*) and the membrane potential (*v*) during AP firing of the left cell (Cell 1). We zoom in on a part of the domain close to the soma and AIS. In this simulation, Cell 1 is stimulated by a somatic stimulation current of *I*^1^_s_ = *−*200 *µ*A/cm^2^. B: Membrane potential in an AIS point of Cell 1 for different values of somatic stimulation, *I*^1^_s_, ranging from 0 to *−*500 *µ*A/cm^2^. C: Membrane potential in an AIS point of Cell 2. The somatic stimulation of Cell 2 is kept fixed at *I*^2^_s_ = *−*27 *µ*A/cm^2^ in all simulations. To quantify the ephaptic effect of Cell 1 stimulation on Cell 2 firing, we measure the distance between the first of the two double spikes of Cell 2 as indicated by the dashed line. In the simulation plotted in Panel C, *I*^1^_s_ = 0, and in Panels A–C the default value of *σ_e_* = 3 mS/cm is applied. D: Cell 2 spike distance as a function of the somatic stimulation strengths of Cell 1 (*I*^1^). We consider the default value of *σ_e_* in addition to the default value reduced by a factor of 5. The spike distance is defined in Panel C.

In Figure 10B, we illustrate the membrane potential at the AIS of Cell 1 for a number of different values of somatic stimulation (*I*^1^) causing AP firing of different frequencies. As we vary the somatic stimulation of Cell 1, the somatic stimulation of Cell 2 is kept constant at *I*^2^ = −27 *µ*A/cm^2^. The resulting Cell 2 AIS membrane potential is plotted in Figure 10C. In that simulation *I*^1^ = 0. However, we want to investigate whether another choice of somatic Cell 1 stimulation, *I*^1^, might alter the spike timing of Cell 2 even though the stimulation of that cell is kept the same. To that end, we define the spike distance for Cell 2 as illustrated in Figure 10C.

In Figure 10D, we report the spike distance of Cell 2 for different Cell 1 stimulation levels. We observe that as the stimulation strength of Cell 1 is increased and Cell 1 fires APs more frequently, the distance between spikes for Cell 2 increases by several milliseconds. We consider the default value of *σ_e_* in addition to a reduced value of *σ_e_/*5. The increase in Cell 2 spike distance as a result of rapid Cell 1 firing is more prominent for the reduced value of *σ_e_*. Since the stimulation of Cell 2 is kept constant and the two cells are not connected by gap junctions or synapses, these spike time alterations must be caused by ephaptic interactions between the neurons.

### 3.5 AP triggering through ephaptic coupling in the presence of high extracellular resistivity

As a final set of examples of ephaptic interactions between neighbouring neurons, we consider the case of an AP being triggered in a neuron due to the EP generated from a nearby firing neuron. We consider the default value of *σ_e_* = 3 mS/cm in addition to three cases of reduced conductivities. The setup used in the simulations is illustrated in Figure 11B.

**Figure 11:**
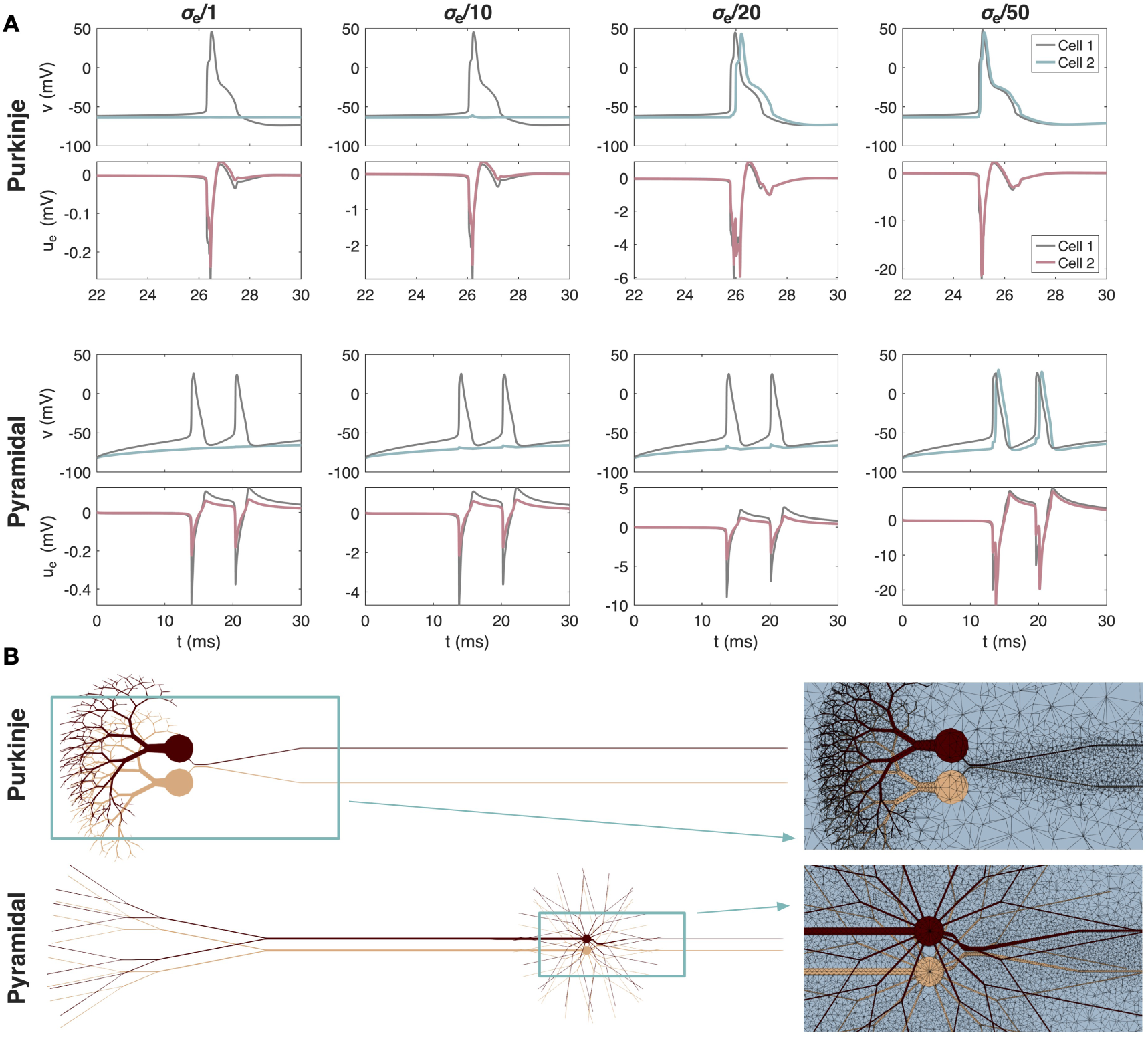
Potential AP triggering through ephaptic coupling between two cerebellar Purkinje neurons and between two neocortical layer 5 pyramidal neurons. A: Membrane potential (*v*) and extracellular potential (*u_e_*) in an AIS membrane point plotted for each cell and four different values of the extracellular conductivity, *σ_e_*. Note that the scaling of the *y*-axis for *u_e_* differs for each value of *σ_e_*. B: Illustration of the simulation setup and a part of the 3D mesh used in the simulations. The AISs of the two neurons are separated by 1 *µ*m of extracellular space. For the Purkinje neurons, Cell 1 is not stimulated and fires spontaneous APs. Cell 2 is made quiescent by a somatic stimulation current of *I*^2^_s_ = 0.7 *µ*A/cm^2^. For the pyramidal neuron, Cell 1 is stimulated by a somatic stimulation current of *I*^1^ = *−*50 *µ*A/cm^2^, causing AP firing, while Cell 2 is stimulated by a weaker somatic stimulation current of *I*^2^_s_ = *−*25 *µ*A/cm^2^, which is not enough in itself to trigger AP firing. For both cell types, the extracellular potential generated during Cell 1 firing is strong enough to trigger AP firing in Cell 2 when *σ_e_* is very small.

For the pyramidal neurons, we let Cell 1 fire APs as a result of a somatic stimulation current of *I*_S_^1^ = −50 *µ*A/cm^2^. Cell 2 is given a weak somatic stimulation current of *I*_S_^2^= −25 *µ*A/cm^2^, which is not strong enough to trigger AP firing in itself. In Figure 11A, we plot the membrane potential and EP in an AIS membrane point of each cell. We observe that for the default value of *σ_e_*, APs are fired in Cell 1, but not in Cell 2. The EP generated during Cell 1 firing reaches a value of about −0.5 mV at the AIS of Cell 1 and about −0.2 mV at the AIS of Cell 2. The corresponding perturbation of the membrane potential, *v* = *u_i_* − *u_e_*, of Cell 2 is not large enough to cause AP firing. As the value of *σ_e_* is reduced, the amplitude of the EP is increased, and we see clear perturbations of the membrane potential of Cell 2. Yet, even for *σ_e_* reduced by a factor of 20, this perturbation is not large enough to trigger an AP. However, when *σ_e_* is reduced by a factor of 50, the membrane potential perturbation of Cell 2 caused by the EP generated during Cell 1 firing is large enough to trigger AP firing at Cell 2 as well.

For the Purkinje neurons, Cell 1 does not receive any somatic stimulation current and fires spontaneous APs, while Cell 2 is given a positive somatic current of *I*^2^ = 0.7 *µ*A/cm^2^, which makes the cell quiescent on its own. However, if the extracellular conductivity is very low (default *σ_e_* divided by a factor of 20 or 50), the negative EP generated from Cell 1 firing is enough to cause firing of Cell 2 as well (see Figure 11A).

## 4 Discussion

This work examines how neurons respond to extracellular stimulation and how local EPs can transmit signals ephaptically from one cell to adjacent cells. Our study is based on a finite element implementation of the EMI model [21, 14, 20], developed using the MFEM finite element software package [33], which allow us to study both endogenous field interactions and responses to imposed stimulation within the same modeling framework. We examine two contrasting cell types – autonomously firing Purkinje neurons and quiescent layer 5 pyramidal neurons. The membrane dynamics of the two neuron types follow [27] and [28], respectively.

The present computations are all performed using a closed-loop (EMI) formulation, in which intra- and extracellular potentials are solved in a fully coupled manner. The more commonly applied open-loop (MI+E) formulation offers computational advantages but falls short in studies of, for example, synchronization of neighboring neurons. In most of the single-neuron examples considered here, an MI+E approach incorporating extracellular stimulation would likely give similar qualitative results as those found here, but previous work [21] and a comparison in the Supplementary Material indicate that quantitative differences may be significant in some parameter regimes, even for a single neuron. However, a systematic assessment of open- vs. closed-loop formulations using realistic geometries and membrane dynamics is beyond the scope of the present paper.

### 4.1 The potential decays as **1*/r*** from a current source

A first step in examining how neurons can influence each other ephaptically is to understand how a single neuron responds to an external current source. This is also of independent interest, as it forms the basis for electrical stimulation techniques used in a range of experimental and clinical settings. The impact of an extracellular current source on a neuron has been investigated experimentally, see, e.g., [7, 29], and the spatial decay of the potential is known analytically from the solution of the electrostatic equation [15]. If *r* denotes the distance in the extracellular space to a point current source, the resulting potential decays as 1*/r*, a pattern illustrated in [7, 29]. The lower panel of Figure 3 follows this decay profile.

### 4.2 The subthreshold membrane potential follows the stimulus frequency

In Figure 4, a weak extracellular current source modulates the subthreshold membrane potential of a nearby neuron, and the resulting oscillation follows the sinusoidal frequency of the stimulus current. The amplitude of the membrane response increases in proportion to the strength of the applied current source. This type of subthreshold frequency-following, with an approximately linear dependence on stimulus amplitude, is consistent with experimental and modeling studies of neurons exposed to weak oscillatory electric fields [7, 29, 39].

### 4.3 Spike frequency is modulated by the strength and distance of a constant current source

In Figure 5 we show how the spike frequency is affected by the strength of an external stimulation source and the distance to this source. For the Purkinje neuron, we first consider the case without somatic stimulation, i.e., *I_s_* ≡ 0, see (1). In this setting, the number of spikes during a 500 ms simulation increases as the extracellular current becomes stronger and as the distance to the source decreases. However, for sufficiently strong stimulation at sufficiently close distances, the AP fails to repolarize properly, and the number of spikes is reduced to a single spike.

Next, we add a positive somatic membrane current to the Purkinje neuron, thereby reducing its excitability. This results in fewer spikes, while the dependence on the strength of the extracellular current source and the distance to the source remains similar to the case without added membrane current. Doubling the additional membrane current continues this trend and yields substantially fewer spikes.

The neocortical layer 5 pyramidal neuron is quiescent in the absence of membrane stimulation (i.e., *I_s_* ≡ 0). In the lower part of Figure 5, we note that for this cell type the extracellular stimulus must be both close and strong in order to initiate spikes. When a sufficiently strong negative *I_s_* is applied to the membrane, spikes are generated, and still more appear when the extracellular stimulus is close and strong.

### 4.4 Spike frequencies are modulated by the frequency of a sinusoidal current source

In Figures 6 and 7 we show how a sinusoidal extracellular current source affects the membrane potential and EP of the two neuronal models. For the Purkinje neuron, a weak low-frequency stimulus has limited impact on the membrane potential, but as the stimulation strength increases the timing of the AP firing is influenced in a phase-dependent manner by the sinusoidal drive. When the phase of the stimulation current produces a sufficiently positive perturbation in the EP, the membrane potential becomes silent and no spikes are generated, whereas the spiking frequency is increased in the opposite phase. These effects are also clear for a 5 Hz stimulation frequency. When the frequency is increased to 30 Hz or 100 Hz, low stimulation amplitudes again have limited impact on the spikes, whereas strong stimulation suppresses spiking during phases of strongly positive EP and activate spiking during phases of negative EP.

In Figure 7, the neocortical layer 5 pyramidal neuron is subjected to a weak somatic membrane stimulation in addition to the sinusoidal extracellular stimulation. We observe that the sinusoidal stimulation induces a combination of occasional spikes and subthreshold modulation rather than regular firing at the drive frequency. Nevertheless, spikes seem to preferentially be generated when the EP is negative corresponding to a negative phase of the stimulation current.

These findings are consistent with experimental studies showing that weak oscillatory electric fields modulate spike timing and entrainment in a cell-type–dependent manner [29] and that transcranial alternating current stimulation (tACS) can phase-lock single-neuron firing *in vivo* [40]. They are also consistent with modeling work based on the Pinsky–Rinzel neuron model [41], which demonstrates changes in firing sensitivity and activity patterns under AC-induced electric fields [42].

### 4.5 Ephaptic interactions between neighboring neurons

In both cardiac electrophysiology and neuroscience, the possibility that one cell can excite its neighbor without direct coupling via gap junctions or chemical synapses has been extensively discussed, see, e.g., [9, 10, 43, 44] for neural tissue and [45, 46, 47] for cardiac tissue. For both organs, this question goes to the core of how tissue organizes excitation waves and is therefore of considerable interest. Here, we assess ephaptic coupling for the two neurons under consideration.

In Figures 8 and 9 we follow up the analysis presented in [14], where we showed that two adjacent cerebellar Purkinje neurons synchronize using the modeling framework applied here. In Figure 8 we illustrate how the APs of two neighboring Purkinje neurons synchronize and how this process depends on the distance between the cells. When the cells are close (1 *µ*m measured at the AIS), the synchronization is rapid and the two APs become completely synchronized. However, when the distance increases to 5 *µ*m, the APs synchronize only partially and reach a stable state in which the spike firing of the two neurons is separated by about 1.2 ms. When the distance is increased to 10 *µ*m, a similar behavior is observed, but the converged distance is about 1.8 ms. The emergence of near-synchronous firing between neighboring Purkinje cells in our simulations is consistent with *in vivo* recordings showing sub-millisecond, tightly synchronized activity of nearby Purkinje cells and population-wide spike alignment [9, 48], and with modeling studies demonstrating that decreasing inter-neuronal spacing or increasing packing density strengthens ephaptic interactions and promotes spike-time synchrony [8, 12].

In Figure 9 we show how the speed of synchronization depends on the value of the extracellular conductivity, *σ_e_*. Panel C demonstrates that synchronization is strongly affected by *σ_e_* in the sense that reducing *σ_e_* markedly increases the speed of synchronization. This is consistent with the observation in [21] that the ephaptic current scales as *O*(1*/σ_e_*). A similar sensitivity of ephaptic effects to the properties of the extracellular space has been reported in computational studies: in networks of ephaptically coupled neurons, stronger ephaptic potentials and reduced inter-neuronal spacing promote spike synchronization and microcluster formation [8], in axon bundles, higher extracellular resistivity (equivalently, lower conductivity) together with increased fibre density enhance ephaptic coupling and facilitate near-synchronous spike volleys [49], while earlier EMI simulations show that the ephaptic current scales approximately as *O*(1*/σ_e_*) and thus becomes negligible for very large extracellular conductivities [21].

In Figure 10 we investigate ephaptic interactions between two neocortical layer 5 pyramidal neurons separated by 1 *µ*m of extracellular space at the AIS. Here, Cell 2 is given a constant weak somatic stimulation current, while the somatic current of Cell 1 is stronger and varied between experiments. Panel A illustrates how an AP firing in Cell 1 perturbs the EP in the narrow cleft between the cells, potentially affecting the dynamics of Cell 2. Indeed, in Panel D, this effect is quantified by reporting the distance between two spikes of Cell 2 as a function of the somatic stimulation of Cell 1. The spike distance is observed to change by several milliseconds depending on the stimulation and resulting firing frequency of Cell 1. Furthermore, the effect increases when *σ_e_* is reduced.

### 4.6 Can a neuron excite its neighbor ephaptically?

As mentioned above, a long-standing question in both cardiac electrophysiology and neuroscience is whether a cell can directly excite its neighbor through the extracellular space alone. We have seen above that one cell can influence another, but can this interaction trigger a full AP? A classical reference is [50], who examined impulse transfer between isolated myocyte pairs. When two single myocytes were gently pushed together in a side-to-side configuration, they observed no electrotonic interaction and no impulse transfer, and concluded that ephaptic transmission was unlikely. This experiment was revisited in [51], but with the cells aligned end-to-end rather than side-by-side. In this geometry, the high-density sodium channel regions at the intercalated disc of the myocytes are brought into proximity, and AP transmission was observed even when gap junctional coupling was absent. Thus, whether signal transfer occurs depends strongly on the cellular configuration.

In Figure 11 we explore this question for our two neuron types placed one *µ*m apart. For the EP generated by one cell to be strong enough to elicit a full AP in its neighbor, the extracellular conductivity *σ_e_* must be sufficiently small. This is consistent with the ephaptic current scaling as *O*(1*/σ_e_*). Whether such small *σ_e_* values occur locally in neural tissue remains uncertain, although geometric confinement and narrow extracellular clefts could, in principle, create domains with reduced effective conductivity. However, in Figure 11, we find that for the EP generated by one cell to directly trigger an AP in a neighboring neuron separated by one *µ*m, the extracellular conductivity, *σ_e_*, must be reduced by roughly a factor of twenty to fifty relative to the standard value of 3 mS*/*cm. We repeated the simulations with the distance between the neurons reduced by one half, but this did not change the conclusion regarding the requirements for direct excitation. This places the *direct triggering* regime outside the range of physiological conductivities. In healthy neural tissue, bulk extracellular conductivity is typically in the range of 3 to 6 mS*/*cm [52].

A notable exception to these bulk values is the Pinceau structure surrounding the Purkinje cell AIS, which forms a locally high-resistivity compartment and supports strong ephaptic inhibition [10]. However, this structure is anatomically and functionally specialized, and its ephaptic effect is inhibitory rather than excitatory. Thus, while Figure 11 shows that direct ephaptic triggering is theoretically possible in the extreme limit of conductivity, it is unlikely to operate as a mechanism for excitatory propagation in typical cortical or cerebellar tissue. Instead, ephaptic effects are more plausibly involved in modulating spike timing and supporting synchronization, as observed at physiological conductivities in Figure 9 and Figure 10.

### 4.7 Limitations

Although the present computational scheme based on the EMI model allows for a more detailed representation of neurons than commonly used cable-based models, there are several important limitations. First, the 3D neuron geometries were generated in Gmsh as idealized morphologies and were not constrained by anatomical reconstructions. Second, the membrane models contain a long list of parameters that are guided by experimental measurements, but these parameters are associated with substantial uncertainty and non-uniqueness, in the sense that quite different parameter sets may reproduce similar activity patterns, [53, 54]. Third, we have assumed that ionic concentrations are constant in space and time within both the intracellular and extracellular domains. These concentrations could have been allowed to vary by, e.g., applying the KNP-EMI model, which combines the EMI framework with an electroneutral electrodiffusion description and allows ion concentrations and reversal potentials to evolve dynamically, [55, 23]. Moreover, at nanometer scales near membranes and in narrow extracellular clefts, the homogenized description used in KNP-EMI breaks down, and electrodiffusive effects, Debye layers and steep local concentration gradients become important. Such effects can be captured only by nanoscale Poisson–Nernst–Planck models, see, e.g., [56, 57, 58], and are not included in the EMI model formulation applied here.

## 5 Conclusion

In this study we used a fully coupled EMI formulation to examine how extracellular stimulation and local field effects influence cerebellar Purkinje neurons and neocortical layer 5 pyramidal neurons. The simulations reproduce the expected 1/*r* decay of potentials from a current source, show that weak fields induce subthreshold oscillations that follow the stimulus frequency, and quantify how constant and sinusoidal stimulation modulate spike rates and timing. For Purkinje neurons, ephaptic interactions synchronize neighboring cells in a distance- and conductivity-dependent manner. In addition, ephaptic interactions between layer 5 pyramidal neurons cause spike time alterations. Together, these results clarify how single-cell biophysics shapes field-mediated interactions.

## Supplementary Information

### Cerebellar Purkinje neuron model parameters

The parameter values for the cerebellar Purkinje neuron are the same as in [14], based on [59]. The expressions for *I*_ion_ and *F* are taken from [59], but intracellular Ca^2+^ dynamics are excluded. The geometry and ion channel conductances of the different parts of the neuron are specified in Tables S1 and S2. In addition, we have used the EMI model parameters *C_m_* = 1 *µ*F/cm^2 [^60], *σ_i_* = 8.2 mS/cm [59], and *σ_e_* = 3 mS/cm [38].

**Table S1:**
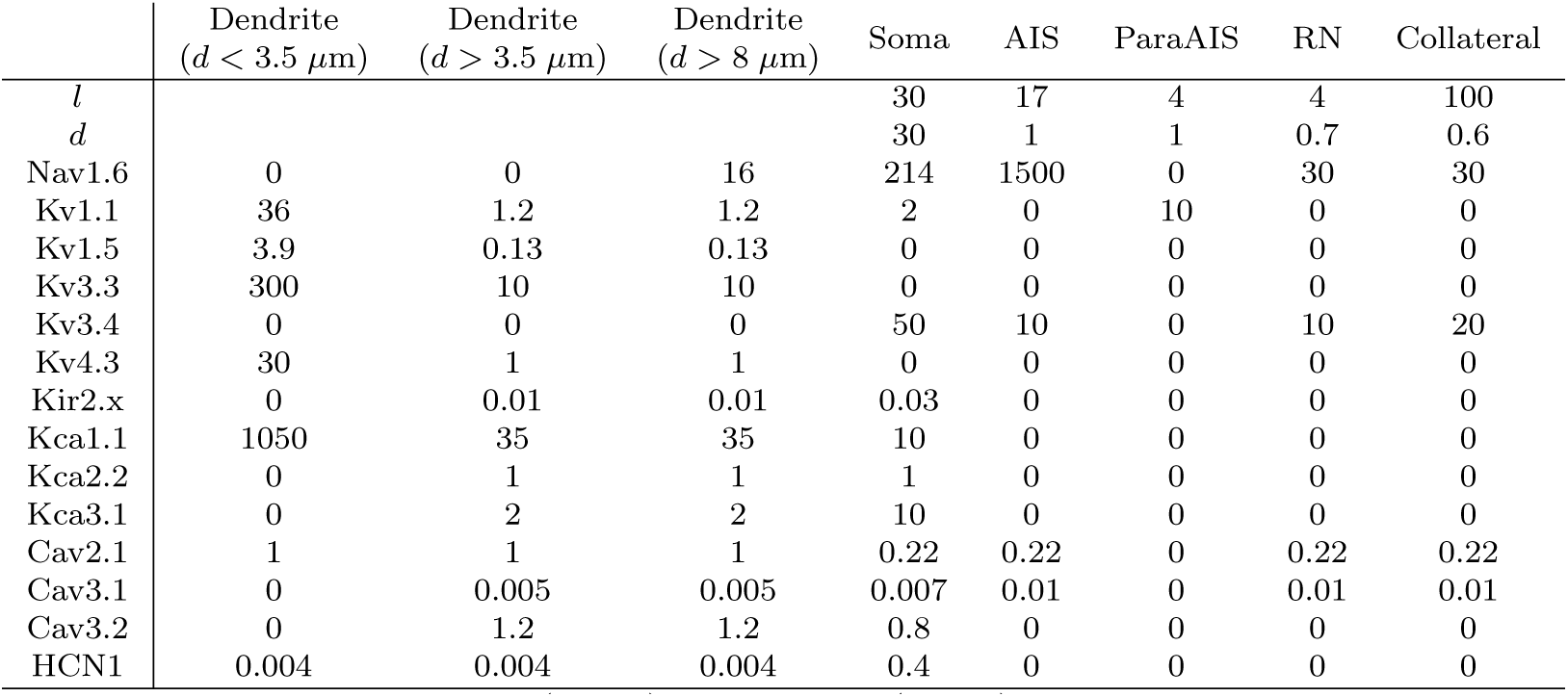
Overview of lengths, *l*, (in *µ*m) diameters, *d*, (in *µ*m) and conductances of different types of ion channels (in mS/cm^2^) in the spatial regions of the Purkinje neuron, based on [59]. There are three Ranvier nodes (RN). Between these, there are myelinated regions of length *l* = 100 *µ*m and diameter *d* = 0.7 *µ*m. For the lengths and diameters of the dendric regions, see Table S2.

**Table S2:**
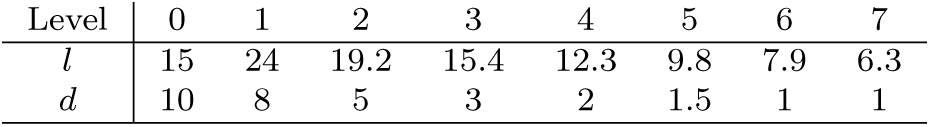
Lengths, *l*, (in *µ*m) and diameters, *d*, (in *µ*m) applied for each branching level in the Purkinje neuron dendrite geometry.

### Neocortical layer 5 pyramidal neuron model parameters

The EMI model representation of a neocortical layer 5 pyramidal neuron model is based the cable model representation in [28], but we have extended the axon modeling applied in [28] to include an axon neck, an axon initial segment (AIS), Ranvier nodes separated by myelinated segments and a collateral. The geometry and membrane model parameters applied in the different neuron segments are specified in Table S3. The geometry is based on [28, 61, 62, 31] and the parameterization is based on the model version from [28] in which the AP is initiated in the axon. The expressions for *I*_ion_ and *F* are taken from [28]. In addition, we have used the EMI model parameters *C_m_* = 1 *µ*F/cm^2 [^60], *σ_i_* = 8.2 mS/cm [28], and *σ_e_* = 3 mS/cm [38].

**Table S3:** Overview of lengths, *l*, (in *µ*m) diameters, *d*, (in *µ*m) and parameters of the Hay et al. membrane model [28] in the spatial regions of the pyramidal neuron. The conductances *g* are given in units of mS/cm^2^, *τ*_decay_ is given in units of ms and, *γ* is unitless. There are three Ranvier nodes (RN). Between these, there are myelinated regions of length *l* = 80 *µ*m and diameter *d* = 1.7 *µ*m. Note that the reported value of *g*_Ca_,_HVA_ and *g*_Ca_,_LVA_ for the apical dendrite correspond to the value in an 200 *µ*m long so-called “hot” zone located about 750 *µ*m to the left of the soma. In the remaining parts of the apical dendrites, the value of *g*_Ca_,_HVA_ is divided by 10 and the value of *g*_Ca_,_LVA_ is divided by 100, following [28].

|  | Apical dendrite | Basal dendrite | Soma | Neck | AIS | RN | Collateral |
| --- | --- | --- | --- | --- | --- | --- | --- |
| $l$ | 1500 | 200 | 25 | 7 | 31 | 4 | 200 |
| $d$ | 1.5–5.9 | 1.0–1.6 | 25 | 3.4 | 3.5 | 1.7 | 1.3 |
| $\bar{g}_{\text{Nat}}$ | 10.745 | 0 | 249.72 | 487.01 | 9740.25 | 487.01 | 487.01 |
| $\bar{g}_{\text{Nap}}$ | 0 | 0 | 0 | 0.292 | 5.834 | 0.292 | 0.292 |
| $\bar{g}_{\text{Kp}}$ | 0 | 0 | 0 | 188.85 | 188.85 | 188.85 | 188.85 |
| $\bar{g}_{\text{Kt}}$ | 0 | 0 | 338 | 77.274 | 77.274 | 77.274 | 772.74 |
| $\bar{g}_{\text{Kv3.1}}$ | 0.904 | 0 | 0 | 473.8 | 473.8 | 473.8 | 473.8 |
| $\bar{g}_{\text{Ca,HVA}}$ | 0.351 | 0 | 0.644 | 0.222 | 0.222 | 0.222 | 0.222 |
| $\bar{g}_{\text{Ca,LVA}}$ | 70.975 | 0.557 | 0.813 | 0.813 | 0.813 | 0.813 | 0.813 |
| $\bar{g}_{\text{SK}}$ | 0.001 | 0 | 99.65 | 0.047 | 0.047 | 0.047 | 0.047 |
| $\bar{g}_{\text{m}}$ | 0.495 | 0 | 0.008 | 13.322 | 13.322 | 13.322 | 13.322 |
| $\bar{g}_{\text{h}}$ | 0.05 | 0.05 | 0.1 | 0.1 | 0.1 | 0.1 | 0.1 |
| $\bar{g}_{\text{leak}}$ | 0.03 | 0.03 | 0.03 | 0.03 | 0.03 | 0.03 | 0.03 |
| $\gamma$ | 0.000637 | 0.000501 | 0.000509 | 0.000525 | 0.000525 | 0.000525 | 0.000525 |
| $\tau_{\text{decay}}$ | 35.7 | 460.0 | 294.7 | 277.3 | 277.3 | 277.3 | 277.3 |

### Comparison of open-loop (MI+E) and closed-loop (EMI) solutions

As discussed in the paper, open-loop algorithms are most frequently used to compute the membrane potential of neurons and the extracellular potential surrounding them [17, 15, 16]. As carefully explained in [16], the open-loop approach involves first solving the membrane problem and then, post hoc, computing the extracellular potential from the membrane currents obtained in the first step. Here, we denote this approach by MI+E, meaning that the membrane (M) equations are solved together with the intracellular (I) space, and the extracellular (E) solution is subsequently computed.

Our aim is now to compare this open-loop approach (MI+E) with the closed-loop EMI model, in which E, M, and I are solved fully coupled. To this end, we define an open-loop version of the EMI model equations. In this approach, the MI system is given by

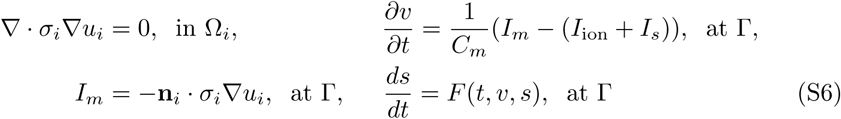

with the assumption *v* = *u_i_* at Γ. After the *u_i_* and *v* solutions are computed by solving the system (S6), *u_e_* is computed by

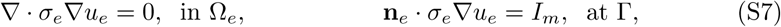

where *I_m_* is collected from the solution of (S6) and the boundary condition on the outer (non-membrane) part of the extracellular boundary is given by *u_e_* = 0. These model definitions allow us to cleanly compare the open-loop and closed-loop approaches in a setup where the open-loop and closed-loop models are as similar as possible, only differing in whether MI is coupled with E or not.

**Figure S1:**
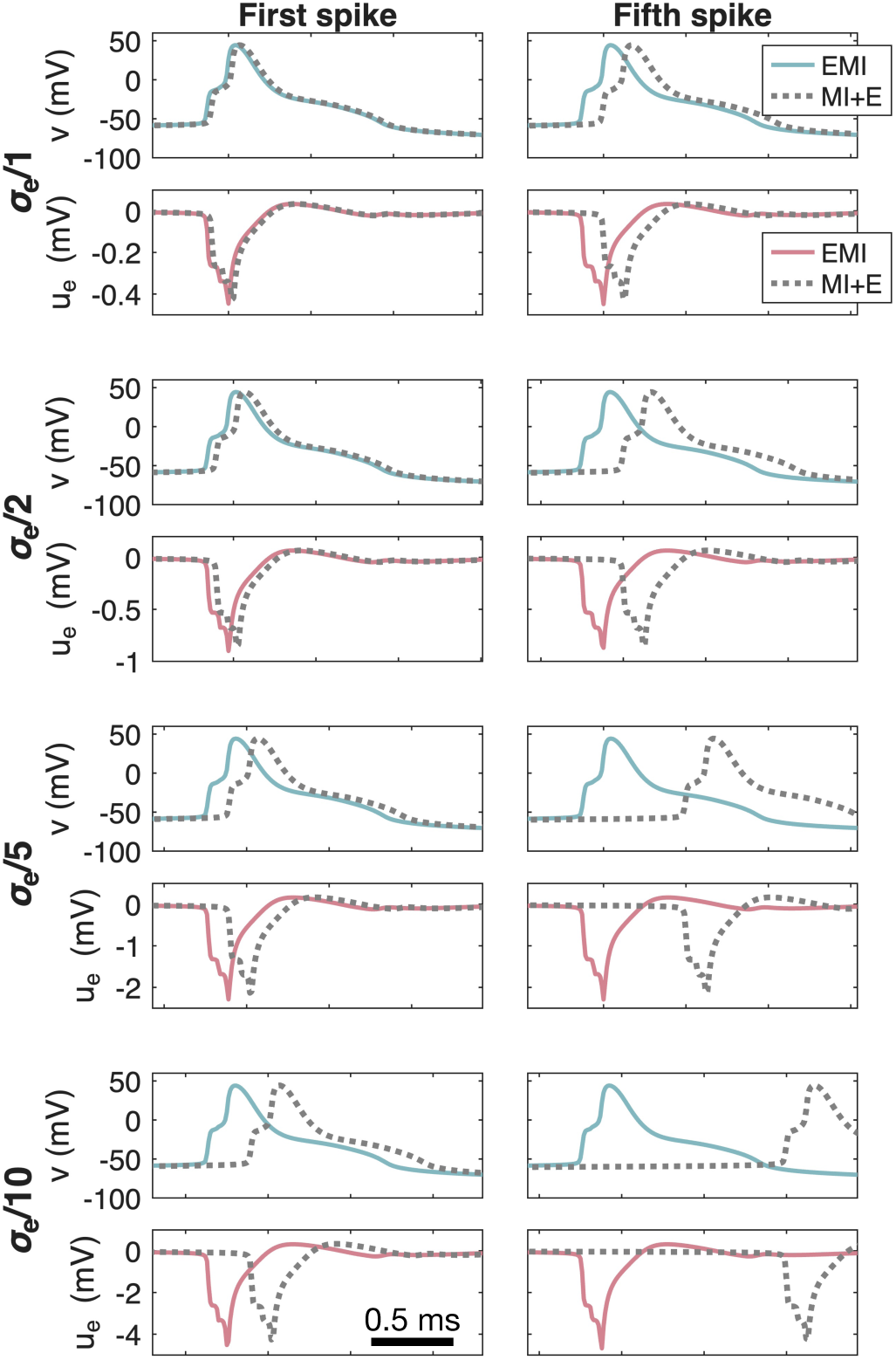
Comparison between open loop (MI+E) and closed loop (EMI) solutions of a spontaneously firing Purkinje neuron. We show the membrane potential, *v*, and the extracellular potential, *u_e_*, in an AIS membrane point during the first and fifth spikes for four different values of the extracellular conductivity, *σ_e_*. No somatic stimulation is applied.

## References

[1] György Buzsáki, Costas A Anastassiou, and Christof Koch. The origin of extracellular fields and currents – EEG, ECoG, LFP and spikes. Nature Reviews Neuroscience, 13(6):407–420, 2012.

[2] Guosong Hong and Charles M Lieber. Novel electrode technologies for neural recordings. Nature Reviews Neuroscience, 20(6):330–345, 2019.

[3] Patrick R Ng, Alan Bush, Matteo Vissani, Cameron C McIntyre, and Robert Mark Richardson. Biophysical principles and computational modeling of deep brain stimulation. Neuromodulation: Technology at the Neural Interface, 27(3):422–439, 2024.

[4] Andres M Lozano, Nir Lipsman, Hagai Bergman, Peter Brown, Stephan Chabardes, Jin Woo Chang, Keith Matthews, Cameron C McIntyre, Thomas E Schlaepfer, Michael Schulder, et al. Deep brain stimulation: current challenges and future directions. Nature Reviews Neurology, 15(3):148–160, 2019.

[5] Joachim K Krauss, Nir Lipsman, Tipu Aziz, Alexandre Boutet, Peter Brown, Jin Woo Chang, Benjamin Davidson, Warren M Grill, Marwan I Hariz, Andreas Horn, et al. Technology of deep brain stimulation: current status and future directions. Nature Reviews Neurology, 17(2):75–87, 2021.

[6] Kristina K Zhang, Rafi Matin, Carolina Gorodetsky, George M Ibrahim, and Flavia Venetucci Gouveia. Systematic review of rodent studies of deep brain stimulation for the treatment of neurological, developmental and neuropsychiatric disorders. Translational Psychiatry, 14(1):186, 2024.

[7] Costas A Anastassiou, Rodrigo Perin, Henry Markram, and Christof Koch. Ephaptic coupling of cortical neurons. Nature Neuroscience, 14(2):217–223, 2011.

[8] R Greg Stacey, Lennart Hilbert, and Thomas Quail. Computational study of synchrony in fields and microclusters of ephaptically coupled neurons. Journal of Neurophysiology, 113(9):3229–3241, 2015.

[9] Kyung-Seok Han, Chong Guo, Christopher H Chen, Laurens Witter, Tomas Osorno, and Wade G Regehr. Ephaptic coupling promotes synchronous firing of cerebellar Purkinje cells. Neuron, 100(3):564–578, 2018.

[10] Antonin Blot and Boris Barbour. Ultra-rapid axon-axon ephaptic inhibition of cerebellar Purkinje cells by the pinceau. Nature Neuroscience, 17(2):289–295, 2014.

[11] Flavio Fröhlich and David A McCormick. Endogenous electric fields may guide neocortical network activity. Neuron, 67(1):129–143, 2010.

[12] Joshua H Goldwyn and John Rinzel. Neuronal coupling by endogenous electric fields: cable theory and applications to coincidence detector neurons in the auditory brain stem. Journal of Neurophysiology, 115(4):2033–2051, 2016.

[13] Gabriel Moreno Cunha, Gilberto Corso, Matheus Phellipe Brasil de Sousa, and Gustavo Zampier dos Santos Lima. Can ephapticity contribute to brain complexity? PloS One, 19(12):e0310640, 2024.

[14] Karoline H Jaeger and Aslak Tveito. Sometimes extracellular recordings fail for good reasons. bioRxiv, pages 2025–07, 2025.

[15] Gary R Holt and Christof Koch. Electrical interactions via the extracellular potential near cell bodies. Journal of Computational Neuroscience, 6(2):169–184, 1999.

[16] Aaron R Shifman and John E Lewis. Elfenn: a generalized platform for modeling ephaptic coupling in spiking neuron models. Frontiers in Neuroinformatics, 13:35, 2019.

[17] Wilfrid Rall. Theory of physiological properties of dendrites. Annals of the New York Academy of Sciences, 96(4):1071–1092, 1962.

[18] Klas H Pettersen, Anna Devor, Istvan Ulbert, Anders M Dale, and Gaute T Einevoll. Current-source density estimation based on inversion of electrostatic forward solution: effects of finite extent of neuronal activity and conductivity discontinuities. Journal of Neuroscience Methods, 154(1):116–133, 2006.

[19] Bartosz Telenczuk, Maria Telenczuk, and Alain Destexhe. A kernel-based method to calculate local field potentials from networks of spiking neurons. Journal of Neuroscience Methods, 344:108871, 2020.

[20] Andres Agudelo-Toro and Andreas Neef. Computationally efficient simulation of electrical activity at cell membranes interacting with self-generated and externally imposed electric fields. Journal of Neural Engineering, 10(2):026019, 2013.

[21] Aslak Tveito, Karoline H Jæger, Glenn T Lines, Łukasz Paszkowski, Joakim Sundnes, Andrew G Edwards, Tuomo Maki-Marttunen, Geir Halnes, and Gaute T Einevoll. An evaluation of the accuracy of classical models for computing the membrane potential and extracellular potential for neurons. Frontiers in Computational Neuroscience, 11:27, 2017.

[22] Alessio P Buccino, Miroslav Kuchta, Karoline H Jæger, Torbjørn Vefferstad Ness, Pierre Berthet, Kent-Andre Mardal, Gert Cauwenberghs, and Aslak Tveito. How does the presence of neural probes affect extracellular potentials? Journal of Neural Engineering, 16(2):026030, 2019.

[23] Ada J Ellingsrud, Andreas Solbrå, Gaute T Einevoll, Geir Halnes, and Marie E Rognes. Finite element simulation of ionic electrodiffusion in cellular geometries. Frontiers in Neuroinformatics, 14:11, 2020.

[24] Karoline H Jæger, Andrew G Edwards, Wayne R Giles, and Aslak Tveito. From millimeters to micrometers; re-introducing myocytes in models of cardiac electrophysiology. Frontiers in Physiology, 12:763584, 2021.

[25] Karoline H Jæger, James D Trotter, Xing Cai, Hermenegild Arevalo, and Aslak Tveito. Evaluating computational efforts and physiological resolution of mathematical models of cardiac tissue. Scientific Reports, 14(1):16954, 2024.

[26] Åshild Telle, James D Trotter, Xing Cai, Henrik Finsberg, Miroslav Kuchta, Joakim Sundnes, and Samuel T Wall. A cell-based framework for modeling cardiac mechanics. Biomechanics and Modeling in Mechanobiology, pages 1–25, 2023.

[27] Stefano Masoli, Diana Sanchez-Ponce, Nora Vrieler, Karin Abu-Haya, Vitaly Lerner, Tal Shahar, Hermina Nedelescu, Martina Francesca Rizza, Ruth Benavides-Piccione, Javier DeFelipe, et al. Human purkinje cells outperform mouse purkinje cells in dendritic complexity and computational capacity. Communications Biology, 7(1):5, 2024.

[28] Etay Hay, Sean Hill, Felix Schürmann, Henry Markram, and Idan Segev. Models of neocortical layer 5b pyramidal cells capturing a wide range of dendritic and perisomatic active properties. PLoS Computational Biology, 7(7):e1002107, 2011.

[29] Soo Yeun Lee, Konstantinos Kozalakis, Fahimeh Baftizadeh, Luke Campagnola, Tim Jarsky, Christof Koch, and Costas A Anastassiou. Cell-class-specific electric field entrainment of neural activity. Neuron, 112(15):2614–2630, 2024.

[30] Karoline H Jæger and Aslak Tveito. Derivation of a cell-based mathematical model of excitable cells. In Modeling Excitable Tissue, pages 1–13. Springer, Cham, 2020.

[31] Douglas J Bakkum, Marie Engelene J Obien, Milos Radivojevic, David Jäckel, Urs Frey, Hirokazu Takahashi, and Andreas Hierlemann. The axon initial segment is the dominant contributor to the neuron’s extracellular electrical potential landscape. Advanced Biosystems, 3(2):1800308, 2019.

[32] Karoline H Jæger, Kristian G Hustad, Xing Cai, and Aslak Tveito. Efficient numerical solution of the EMI model representing the extracellular space (E), cell membrane (M) and intracellular space (I) of a collection of cardiac cells. Frontiers in Physics, 8:539, 2021.

[33] Robert Anderson, Julian Andrej, Andrew Barker, Jamie Bramwell, Jean-Sylvain Camier, Jakub Cerveny, Veselin Dobrev, Yohann Dudouit, Aaron Fisher, Tzanio Kolev, Will Pazner, Mark Stowell, Vladimir Tomov, Ido Akkerman, Johann Dahm, David Medina, and Stefano Zampini. MFEM: A modular finite element methods library. Computers & Mathematics with Applications, 81:42–74, 2021.

[34] MFEM: Modular finite element methods [Software]. mfem.org.

[35] Christoph Geuzaine and Jean-François Remacle. Gmsh: a three-dimensional finite element mesh generator with built-in pre- and post-processing facilities. International Journal for Numerical Methods in Engineering, 79:1309–1331, 2009.

[36] Stanley Rush and Hugh Larsen. A practical algorithm for solving dynamic membrane equations. IEEE Transactions on Biomedical Engineering, 4:389–392, 1978.

[37] Johan Hake, Henrik Finsberg, Kristian Gregorius Hustad, and George Bahij. Gotran – General ODE TRANslator, 2020. https://github.com/ComputationalPhysiology/gotran.

[38] Gaute T Einevoll, Christoph Kayser, Nikos K Logothetis, and Stefano Panzeri. Modelling and analysis of local field potentials for studying the function of cortical circuits. Nature Reviews Neuroscience, 14(11):770–785, 2013.

[39] Jacqueline K Deans, Andrew D Powell, and John GR Jefferys. Sensitivity of coherent oscillations in rat hippocampus to ac electric fields. The Journal of physiology, 583(2):555–565, 2007.

[40] Matthew R Krause, Pedro G Vieira, Bennett A Csorba, Praveen K Pilly, and Christopher C Pack. Transcranial alternating current stimulation entrains single-neuron activity in the primate brain. Proceedings of the National Academy of Sciences, 116(12):5747–5755, 2019.

[41] Paul F Pinsky and John Rinzel. Intrinsic and network rhythmogenesis in a reduced traub model for ca3 neurons. Journal of Computational Neuroscience, 1(1):39–60, 1994.

[42] Chunhua Yuan, Rupei Chen, Xiangyu Li, and Yueyang Zhao. Effects of ac induced electric fields on neuronal firing sensitivity and activity patterns. Frontiers in Computational Neuroscience, 19:1612314, 2025.

[43] Costas A Anastassiou and Christof Koch. Ephaptic coupling to endogenous electric field activity: why bother? Current Opinion in Neurobiology, 31:95–103, 2015.

[44] Hemant Bokil, Nora Laaris, Karen Blinder, Mathew Ennis, and Asaf Keller. Ephaptic interactions in the mammalian olfactory system. J Neurosci, 21:1–5, 2001.

[45] Rengasayee Veeraraghavan, Robert G Gourdie, and Steven Poelzing. Mechanisms of cardiac conduction: a history of revisions. American Journal of Physiology-Heart and Circulatory Physiology, 306(5):H619–H627, 2014.

[46] Robert G Gourdie. The cardiac gap junction has discrete functions in electrotonic and ephaptic coupling. The Anatomical Record, 302(1):93–100, 2019.

[47] Karoline H Jæger, Ena Ivanovic, Jan P Kucera, and Aslak Tveito. Nano-scale solution of the Poisson-Nernst-Planck (PNP) equations in a fraction of two neighboring cells reveals the magnitude of intercellular electrochemical waves. PLoS Computational Biology, 19(2):e1010895, 2023.

[48] Ehsan Sedaghat-Nejad, Jay S Pi, Paul Hage, Mohammad Amin Fakharian, and Reza Shadmehr. Synchronous spiking of cerebellar purkinje cells during control of movements. Proceedings of the National Academy of Sciences, 119(14):e2118954119, 2022.

[49] Helmut Schmidt and Thomas R. Knösche. Modelling the effect of ephaptic coupling on spike propagation in peripheral nerve fibres. Biological Cybernetics, 116(4):461–473, 2022.

[50] R Weingart and P Maurer. Action potential transfer in cell pairs isolated from adult rat and guinea pig ventricles. Circulation Research, 63(1):72–80, 1988.

[51] Jens Waaben, Johannes P Hofgaard, Greta L Pick, Christopher Loisel, Thomas Engstrom, Niels-Henrik Holstein-Rathlou, Steven Poelzing, and Morten Schak Nielsen. Ephaptic coupling enables action potential conduction. bioRxiv, pages 2024–11, 2024.

[52] Nikos K Logothetis, Christoph Kayser, and Axel Oeltermann. In vivo measurement of cortical impedance spectrum in monkeys: implications for signal propagation. Neuron, 55(5):809–823, 2007.

[53] Eve Marder and Adam L Taylor. Multiple models to capture the variability in biological neurons and networks. Nature Neuroscience, 14(2):133, 2011.

[54] Astrid A Prinz, Dirk Bucher, and Eve Marder. Similar network activity from disparate circuit parameters. Nature Neuroscience, 7(12):1345–1352, 2004.

[55] Ada J Ellingsrud, Cécile Daversin-Catty, and Marie E Rognes. A cell-based model for ionic electrodiffusion in excitable tissue. In Modeling Excitable Tissue, pages 14–27. Springer, Cham, 2021.

[56] Jurgis Pods. A comparison of computational models for the extracellular potential of neurons. Journal of Integrative Neuroscience, 16(1):19–32, 2017.

[57] Jurgis Pods, Johannes Schönke, and Peter Bastian. Electrodiffusion models of neurons and extracellular space using the Poisson-Nernst-Planck equations – numerical simulation of the intra- and extracellular potential for an axon model. Biophysical Journal, 105(1):242–254, 2013.

[58] Karoline H Jæger and Aslak Tveito. Electrodiffusion dynamics in the cardiomyocyte dyad at nano-scale resolution using the Poisson-Nernst-Planck (PNP) equations. PLOS Computational Biology, 21(6):e1013149, 2025.

[59] Stefano Masoli, Sergio Solinas, and Egidio D’Angelo. Action potential processing in a detailed purkinje cell model reveals a critical role for axonal compartmentalization. Frontiers in Cellular Neuroscience, 9:47, 2015.

[60] Silvio Weidmann. Electrical constants of trabecular muscle from mammalian heart. The Journal of Physiology, 210(4):1041–1054, 1970.

[61] Sandrine Romand, Yun Wang, Maria Toledo-Rodriguez, and Henry Markram. Morphological development of thick-tufted layer V pyramidal cells in the rat somatosensory cortex. Frontiers in Neuroanatomy, 5:5, 2011.

[62] Patrik Krieger, Christiaan PJ de Kock, and Andreas Frick. Calcium dynamics in basal dendrites of layer 5A and 5B pyramidal neurons is tuned to the cell-type specific physiological action potential discharge. Frontiers in Cellular Neuroscience, 11:194, 2017.

